# Two Separated Worlds: on the Preference of Influence in Life Science and Biomedical Research

**DOI:** 10.1101/2024.05.10.592442

**Authors:** Zuguang Gu

## Abstract

We introduced a new metric, “citation enrichment”, to measure country-to-country influence using citation data. This metric evaluates the degree to which a country prefers to cite another country compared to a random citation process. We applied the citation enrichment method to over 12 million publications in the life science and biomedical fields and we have the following key findings: 1) The global scientific landscape is divided into two separated worlds where developed Western countries exhibit an overall mutual under-influence with the rest of the world; 2) Within each world, countries form clusters based on their mutual citation preferences, with these groupings strongly associated with their geographical and cultural proximity; 3) The two worlds exhibit distinct patterns of the influence balance among countries, revealing underlying mechanisms that drive influence dynamics. We have constructed a comprehensive world map of scientific influence which greatly enhances the deep understanding of the international exchange of scientific knowledge. The citation enrichment metric is developed under a well-defined statistical framework and has the potential to be extended into a versatile and powerful tool for bibliometrics and related research fields.

## 1. Introduction

Scientific publications serve as the primary medium for storing and disseminating scientific knowledge. They play a vital role in documenting research findings, advancements, and insights across various fields of research. These publications encompass a wide range of formats, including research articles, reviews, conference papers, reports and scientific books. Among them, knowledge flows are mainly represented as citations between publications where researchers use them as an acknowledgement of prior research, utilization of established methods and tools, and validation for their scientific findings. In short, citations can be used as a proxy for measuring scientific influence (Aksnes et al., 2019). In various evaluation systems, citation has been frequently used as an important indicator for ranking publications, authors, journals or institutions (Roldan-Valadez et al., 2019). However, despite its simplicity and popularity, citations have been criticized for its oversimplification and inappropriateness in measuring the influence of scientific outcomes, including the bias towards established researchers and high-impact journals (Tahamtan and Bornmann, 2018; Urlings et al., 2021), or overestimation from self-citations and citation manipulation (Wren and Georgescu, 2022). What’s more, even in an individual publication, the cited publications do not influence the citing publication equally (Zhang and Wu, 2021). Nevertheless, on the macro-level, e.g., when evaluating the scientific influence of countries, citations remain one of the most widely used metrics for studying global patterns and dynamics of scientific knowledge exchange (Tahamtan and Bornmann, 2019).

A wide range of bibliometric methods have been developed to measure a country’s scientific influence using citation data (Radicchi and Castellano, 2015). Among the most commonly used approaches are count-or fraction-based metrics, such as the total number or the fraction of citations a country contributes or receives from the globe (Donthu et al., 2021). More complex metrics have also been proposed. For example, for a given country denoted as *A* and the set of all international publications not published in *A* as *P*, Hassan and Haddawy (2013) measured the international impact of *A* as the fraction of citations from *P* that cite *A*, relative to the total citations from *P*. This metric represents the likelihood of *A* being cited by an international publication. However, it tends to generate very small values for countries with lower citation volumes because the total number of citations from *P* is usually huge, which makes the interpretation of the impact values difficult. Carammia (2022) assessed the country-level international influence in the field of political science by counting papers published in high-impact journals, which are assumed to be internationally recognized. The limit of this study is, there was no quantitative measurement of the internationalization of these journals and the relation of the internationalization between journals and papers still needs more careful explorations. Lancho Barrantes et al. (2012) proposed a metric named “impact on papers per paper” by taking the citation network as a bipartite graph where cited papers and citing papers are the two distinct sets^1^, then the metric is defined as the number of links in the graph divided by the number of cited and citing papers from a given country, i.e., the average number of citations per cited paper and per citing paper. Apparently, if the bipartite graph is fully connected, the metric has a value of 1, but in real cases, the bipartile citation graphs are sparse, resulting in very small metric values that are not always intuitive to interpret.

For a country *A*, most existing studies summarize its scientific influence by aggregating citation data from its citing countries into a single metric (Bogart et al., 2021). These approaches provide a straightforward way for comparing and ranking countries and studying longitudinal trends in country-level influence (Abramo et al., 2019; Carammia, 2022; Hassan and Haddawy, 2013; Lancho Barrantes et al., 2012), but it has obvious limitations. First, the distribution of scientific activity is highly uneven across the globe, with the majority volumes of scientific knowledge coming from a small group of science-advanced countries (Elango and Oh, 2022). Simply averaging all citations of *A* without accounting for country-specific differences of *A*’s citing countries neglects the effects from “scientifically small” countries. Second, the flow of scientific knowledge can vary significantly across regions due to political, social, and cultural differences (Quatraro and Usai, 2017). Aggregating citations into a single value makes it impossible to study the interesting region-specific science dissemination patterns from *A*’s citing countries. To reveal such details missing in current studies, it is necessary to extend the country-wise analysis into two dimensions where a country-to-country influence analysis is conducted. Unfortunately, very few studies have explored this direction. Abramo et al. (2019) measured the balance of a country’s knowledge flow as the difference between its international citations and international references. While the focus of that study was still to assign a general “balance value” country-wise, it could be adapted for country-to-country analysis by only looking at a single second country instead of merging all international countries. However, there were only four countries investigated in the study which is insufficient for discovering broader patterns on the global level.

At the macro-level, the scientific influence of country *B* on country *A* can be evaluated by the total number of citations *B* receives from *A* or the fraction of these citations to the total citations made by *A*. Such methods often rank science-advanced countries with high citation volumes, in most cases the US, at the top of *A*’s most influential countries list (Carammia, 2022). Indeed, countries with large science outputs have major influences on the citing countries globally. For instance, for all the countries investigated in this study, the US is their most cited country with an average of 20.4% of their total citations. However, this raises two questions. First, note the number of publications in the US is also dramatically high (28.5% of global publications), suggesting that the high influence from the US is basically due to the accumulation of the massive publications produced in it. At the individual publication level, this prompts the first question: Given a single publication from the US, how likely does *A* cite it? Second, as the unequal distribution of publications among countries, assume a publication in *A* searches references totally randomly to cite, on average it cites the US much more frequently than other countries. This means the US has a high influence on *A* even if the citing preference of *A* is “unconscious”. This raises the second question: How much of *A*’s citations to *B* are driven by a “subjective intention” to cite *B*, as opposed to an “unconscious” preference arising from random citations?

Most current studies on country-level scientific influence tend to highlight science-advanced countries (Cancino et al., 2017; Wondimagegn et al., 2023), while there are very few studies that reveal the influence patterns of science-small countries at the global scale. This may lead to a biased and incomplete view of global science dissemination. For example, consider Norway and Finland, both as science-small countries. Norway ranks 25th and Finland ranks 22nd in terms of their total citations received from the globe. Their science volumes are too small that they are not often selected in the global studies on scientific influences, while only discussed in regional studies specifically limited on Northern Europe (Narayan et al., 2024). Specifically, only 0.92% of Finland’s references are from Norway with a rank of 18, and 1.2% of Norway’s references are from Finland with a rank of 16. It is really difficult to conclude Finland and Norway have a strong mutual influence from these statistics. However, it is almost common sense that Norway and Finland are highly related both geographically and historically, and they have strong collaborations across their economics, culture, and science (Liimatainen, 2023). It should be expected that they exhibit a strong preference to influence each other that existing metrics fail to capture. This highlights the need for a new metric capable of accurately measuring the preference of scientific influence between countries, independent of their overall citation volumes. Such a metric would provide a more equitable perspective on global scientific influence, particularly for underrepresented science-small countries.

In this work, we proposed a new country-to-country metric named “*citation enrichment*” which aims to address the questions we have raised. Being different from other metrics used in current studies, the citation enrichment metric evaluates scientific influence from a new perspective by answering the question: “Does a country cite another country more preferably?” by comparing it to a random citation process. Methodologically, citation enrichment has two advantages. First, it is a relative metric of which all countries are against the same baseline where publications cite randomly. Second, the random model considers the citation volumes of countries and the metric corrects for the effects from larger countries that have higher probabilities of being cited. This metric captures both over-influence (when a country is cited more frequently than expected under random citation) and under-influence (when a country is cited less frequently than expected). Additionally, the enrichment metric has a well-defined form for theoretical statistical analysis and it has already been widely applied in other fields such as biological data analysis (Rivals et al., 2006).

The citation enrichment metric corrects for the science volumes of countries, thus it provides a robust and standardized way for studying mutual influence patterns among all country pairs from the same scale. To demonstrate the meaningfulness and applicability of the citation enrichment metric in revealing global patterns of scientific influence, we performed an exploratory and empirical analysis on the PubMed dataset of more than 12 million publications from 2000 to 2023 in the life science and biomedical fields. Note, that the aim of this empirical study is not to rank individual countries but to provide a comprehensive world map of international scientific influences between countries. We have the following findings from the analysis. 1. Science-developed countries have a clear isolation from science-developing countries, where the preference of their mutual citations is weaker than random. 2. The 72 countries involved in this analysis can be partitioned into seven distinct groups based on their mutual citation enrichment patterns, with these groups strongly associated with their geographical classifications and cultural proximity. 3. Citation balance analysis reveals mechanisms underlying the dynamic citation processes between countries and the global scientific environment.

In this article, we introduced the citation enrichment metric and explored its empirical applications from a data science perspective. The remainder of this article is organized as follows. In Section 2, we described the identification of domestic publications from the PubMed dataset. In Section 3, we formally defined the citation enrichment metric, as well as providing its interpretation from the probabilistic perspective. Next in the Results section, we presented the empirical analysis on the PubMed dataset. We started with a traditional fraction-based citation analysis in Section 4.1 to demonstrate its limitation of how it generates insufficient and inappropriate conclusions. From Section 4.2 we performed the citation enrichment analysis both on domestic and international citations. We first revealed that domestic citations exhibit a significantly stronger preference compared to international citations in Section 4.2, accompanied by a theoretical interpretation. Then in Section 4.3, we constructed a comprehensive world map of the preference of science influences between countries. We performed a clustering analysis and demonstrated that country groupings have strong associations with their geographical locations and cultural similarities. As the citation is represented as a directional relation, in Section 4.4 we further studied the balance of citation preference of a country to the globe. We revealed different dynamics of citation preference across different regions. Last, in Section 5, we discussed the implications and limitations of the citation enrichment metric. In particular, we discussed in depth the extended studies that can be derived from the citation enrichment analysis, from the methodological perspective.

## 2. Data and preprocessing

### 2.1. Data source

PubMed is the world’s largest public database of literature for biomedical and life science research. We downloaded XML files from the PubMed FTP^2^ with file indices from 1 to 1366 (by 2024-04-07). Publications between the years 2000 and 2023, written in English, and with numbers of authors between 1 and 50 were included. 21,366,669 publications were used as the starting dataset of this study.

### 2.2. Country name identification

The identification of country names from affiliation texts is dictionary-based. In many bibliometrics studies, countries were directly matched from affiliation texts by a list of pre-compiled country names, such as in the tool *bioliometrix* (Aria and Cuccurullo, 2017). This approach requires affiliations to be well-formatted. It can almost be ensured in recent publications in which journals have stringent formatting requirements for electronic archiving, but affiliations are more loosely formatted in earlier publications. Country names may have aliases, e.g., United States of America/United States/USA/US or Germany/Deutschland, thus, an incomplete list of country names may cause countries to fail to be identified. It also occurs often that country names are not even encoded in affiliations. For example, a large number of publications from the US only encode affiliations at the state level. Therefore, we built complex dictionaries that contain information on various levels for constructing more comprehensive mappings between affiliations and countries, which include names of countries^3^, cities^4^, institutions, states/provinces^5^, and email domain names. We also manually collected 689 mappings on various levels to official country names. The scripts for identifying country names are available at https://github.com/jokergoo/citation_analysis.

98.1% of all affiliations were mapped to single countries. We manually validated the identified countries by randomly selecting 1,000 affiliation texts. All 1,000 affiliations were correctly identified, thus we estimate the misidentification rate to be less than 0.1%.

### 2.3. Domestic and international publications

Publications can be categorized into two groups: domestic publications where all authors are from the same country, and international publications where the studies are conducted by authors from multiple countries (Figure 1). Domestic publications represent scientific knowledge generated within individual countries, while international publications reflect knowledge shared across countries. Citations are directional and link publications within and between both groups. In international publications, it is hard to associate them with a single country, making the source country of the scientific knowledge ambiguous. On the other hand, globally, domestic publications are the major type of publications (Chinchilla-Rodríguez et al., 2019; Lancho Barrantes et al., 2012). In the PubMed dataset used for this study, the number of domestic publications is 7.7 times greater than the number of international publications. Therefore, in this study, we only focused on domestic publications and the citation flows among them (green box in Figure 1).

**Figure 1.**
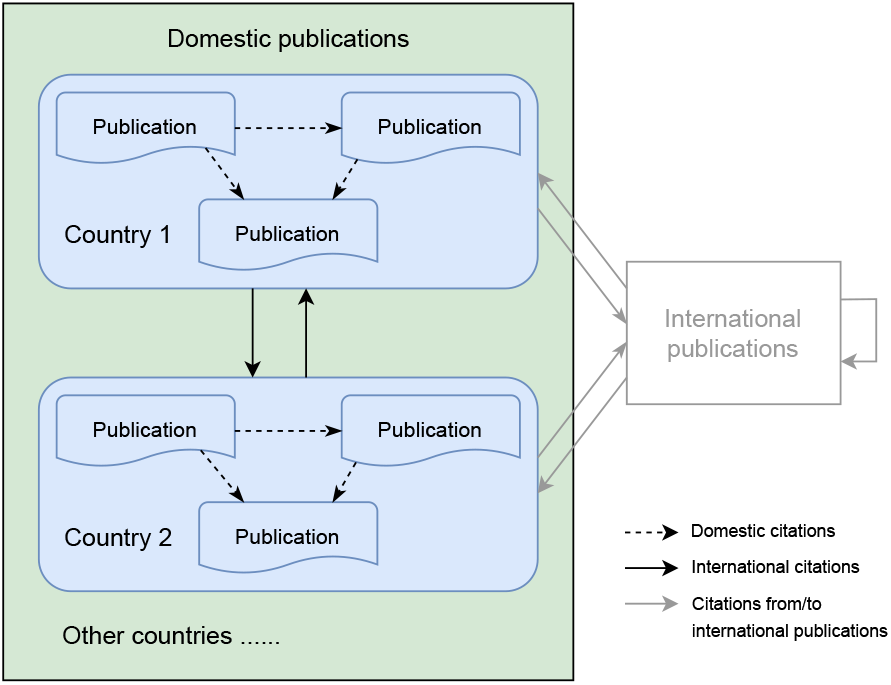
Categories of scientific publications and citations. Domestic publications: papers where authors are from the same country; International publications: papers where authors are from multiple countries. Arrows represent directions of citations. International publications and their citation relations are colored in grey because they are not included in this study.

After countries were identified from authors’ affiliations, in a publication, each author is associated with no country (if the author has no affiliation information or the country can not be identified), a single country or multiple countries. Domestic publications were identified based on the following rules:

1. We assume the last author is the most senior in the publication and should be associated with exactly one country if the publication is domestic.

2. The first author is allowed to have no country associated, or the first author should only have one associated country which should be the same as the country associated with the last author.

3. If there are more than two authors, the publication is identified as domestic if no less than 80% of authors are associated with the same country.

In this study, we additionally restricted our analysis to the publications only published in the Web of Science (WoS) core journals collection^6^. We also removed isolated publications that do not cite or are not cited by any other publication in the dataset. Finally, there were 139,760,456 citations from 12,033,156 domestic publications, with each citation linking two domestic publications.

Please note, in current studies, it is a common practice that a domestic publication strictly requires all authors to be affiliated with the same country. Domestic publication is a measurement of the scientific knowledge produced within a country. In this study, we also included publications we term “approximately domestic publications” in rule 3, which allows for a very small proportion (less than 20%) of international authors to be listed in the middle of the author list. This rule is based on several considerations from our experience as researchers in the field of biomedicine and life science. 1. Authors listed in the middle of the author list generally contribute less significantly to the scientific research presented in the publication. 2 Research in biomedicine and life sciences often spans long periods, during which researchers may change affiliations. 3. Short-term research positions, such as visiting scholars from other countries, can lead to international affiliations on author lists. Although these factors may make the author list partially international, we still classify these publications as domestic because we believe the core scientific knowledge is produced domestically within the country. In this study, approximately domestic publications account for 3.9% of all domestic publications (more comparisons on approximately domestic publications can be found in Supplementary File 1).

## 3. Methods

### 3.1. Motivation

Assume publications from country *A* cite a total of *n* times where they cite publications from country *B* for *k* times, then it is natural to use the fraction of *A* citing *B* to the total number of citations from *A*, i.e., *k/n*, as the metric of how *B* influences *A*. As an example in Figure 2A, for all citations made by mainland China, 30.4% of them reference the US, which is the largest source of citing countries of mainland China, even larger than mainland China citing itself (25.6%). As a comparison, only 0.52% of citations from mainland China reference Singapore. Indeed it is reasonable to conclude that the US has a dominantly stronger influence than Singapore on mainland China by directly comparing the magnitude of the citing fractions. However, notice that the amounts of publications of the US and Singapore differ significantly which leads to the total volumes of global citations to the US and Singapore also differing significantly (59.2M vs. 0.53M). This in turn results in, 42.4% of the global citations referencing the US and only 0.38% of them referencing Singapore. Let’s consider a random scenario where a publication randomly picks another publication to cite, and then naturally countries with larger citation volumes have higher probabilities of being cited. Country-wise, taking the US as an example, the global citing fraction can be used as an estimation of the probability (*p* = 0.424) of the US being cited in the random citation model. Then, if there is no preference for citing individual publications from the US, all the countries should have an equal probability of *p* = 0.424 citing the US. Thus, we can measure the preference of mainland China citing the US as the fold enrichment of the observed citing fraction to the expected citing fraction, i.e., 30.4%/42.4% = 0.72 (Figure 2B). The fold enrichment is less than one, and under this context, we can conclude mainland China is under-influenced by the US where it cites the US less frequently than random. As a comparison, the fold enrichment of mainland China citing Singapore is 0.52%/0.38% = 1.37 which is larger than one, meaning mainland China is over-influenced by Singapore. Overall, we can conclude mainland China has a stronger preference of being influenced by Singapore than the US.

**Figure 2.**
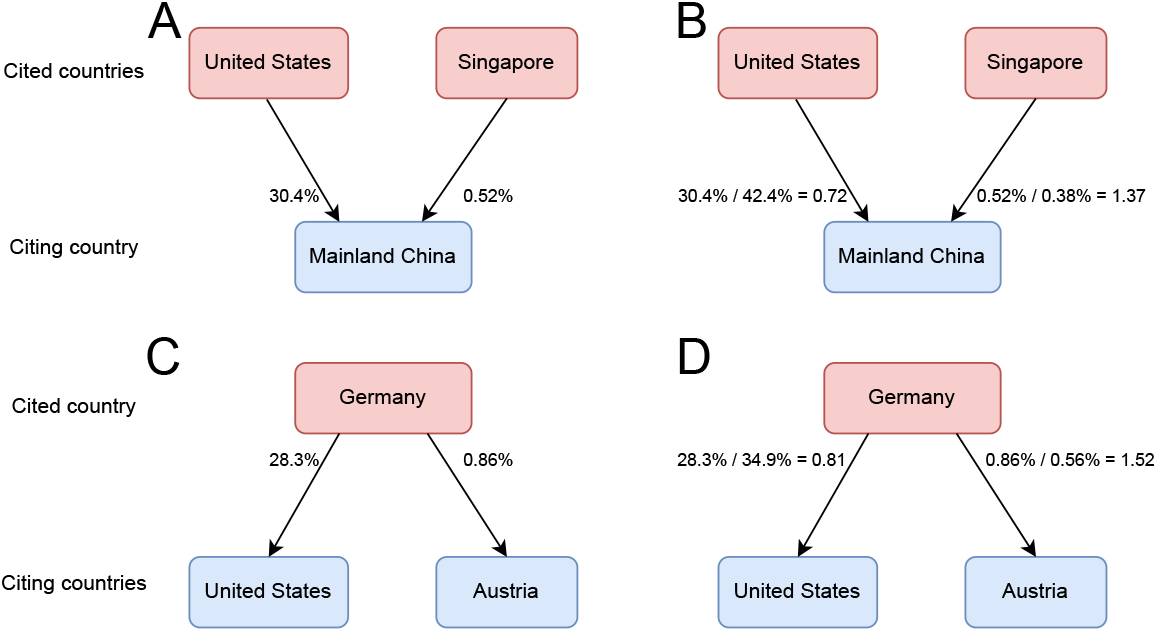
Compare the fraction-based method and the fold enrichment method for measuring the preference of scientific influence from citation data. A) 30.4% and 0.52% of the citations made by mainland China reference the US and Singapore. B) The fold enrichment of mainland China citing the US and Singapore are 0.72 and 1.37. C) 28.3% and 0.86% of Germany’s citations are contributed by the US and Austria. D) The fold enrichment of Germany being cited by the US and Austria are 0.81 and 1.52.

It is similar if we look at the citation enrichment of how a country is cited, i.e., how it distributes its influence to other countries. As an example in Figure 2C, for all citations that Germany receives, 28.3% of them are made by the US, which is the largest country that cites Germany, even larger than Germany cited by itself (17.6%). However, 34.9% of the global citations are made by the US, which makes the fold enrichment of Germany cited by the US to be 28.3%/34.9% = 0.81, meaning Germany has an under-influence on the US (Figure 2D). As an opposite example, 0.86% of Germany’s citations and 0.56% of global citations are made by Austria, generating a fold enrichment of 0.86%/0.56% = 1.52, implying that Germany has an over-influence on Austria (Figure 2D).

For both examples, the citation enrichment highlights the tight and positive relations between Singapore and mainland China, as well as between Germany and Austria, although their absolute citation volumes differ greatly. These results suggest a strong association between the citation preference and the closeness in the economics, culture, and science in these country pairs.

### 3.2. Citation enrichment

Formally, to calculate the fold enrichment of how a country *A* cites country *B* or how *B* is cited by *A*, let’s denote the total number of global citations as *N*, the total number of citations made from *A* as *n*, the total number of citations that *B* receives as *m*, the number of citations of *A* citing *B* as *k*, then the number of citations in different categories can be written into the 2 × 2 contingency table in Table 1.

**Table 1:** The 2 × 2 contingency table of the numbers of citations.

| | Country $B$ (cited) | Other countries (cited) | Total |
| --- | --- | --- | --- |
| Country $A$ (citing) | $k$ | $n - k$ | $n$ |
| Other countries (citing) | $m - k$ | $N + k - m - n$ | $N - n$ |
| Total | $m$ | $N - m$ | $N$ |

The definition of fold enrichment can be given in three different ways but generates identical values, reflecting different aspects of interpreting the enrichment. First, if we read Table 1 horizontally, the fold enrichment denoted as *r* can be defined as the ratio of the following two fractions:

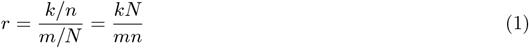

where *k/n* is the fraction of *B* being cited only in *A*-citing publications, and *m/N* is the global fraction of *B* being cited. When *r >* 1, it means that on average *B* is cited by *A* more than *B* is cited globally. Thus, there is an over-representation of *B* being cited by *A* compared to the global average. We can conclude that *B* has an over-influence on *A*. And when *r* < 1, we conclude that *B* has an under-influence on *A*.

Second, if we read Table 1 vertically, *r* can be defined as the ratio of two fractions in a different form:

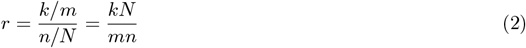

where *k/m* is the fraction of citations from *A* only in *B*-cited publications, and *n/N* is the global fraction of citations from *A*. When *r >* 1, it means that on average *A* cites *B* more frequently than *A* cites globally. Thus, there is an over-representation of *A* citing *B*. We conclude that *A* has an over-influence from *B*.

Third, we can read Table 1 in both dimensions in a neutral way. We first calculate the following two probability values denoted as *p*_1_ and *p*_2_:

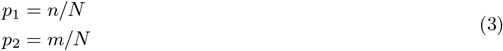

where *p*_1_ is the probability of a citation from *A* (i.e., cited by *A*) and *p*_2_ is the probability of *B* being cited in the globe. If the two events of *A* citing and *B* being cited are independent, we can multiply *p*_1_ and *p*_2_ to obtain an expected number of citations from *A* to *B*, denoted as *k*_exp_:

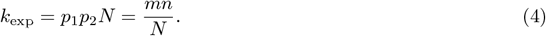

Then the fold enrichment *r* is defined as the ratio of the observed number of citations to the expected number:

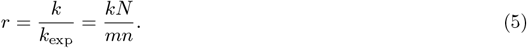

It is straightforward to see that the three definitions of fold enrichment *r* generate the same value. In this study, log2-transformation is additionally applied to the fold enrichment to generate the final metric of the enrichment score, denoted as *s*:

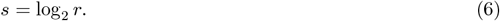

In this article, we say that *B* has an over-influence on *A* or *A* has an over-influence from *B* if *s* > 0, and *B* has an under-influence on *A* or *A* has an under-influence from *B* if *s* < 0. If it is necessary to clarify the direction of the influence, we will explicitly add subscripts to the notations such as *k*_*B*-*A*_, *r*_*B*-*A*_, *s*_*B*-*A*_, *m*_*B*-_, and *n*_-*A*_ where the first letter before “-” corresponds to the cited country and the second letter corresponds to the citing country. If the letter is empty, it means the globe. If *A* is the same as *B*, the enrichment score, i.e., *s*_*A*-*A*_, measures the domestic preference of influence of the country.

The citation network can be constructed as a bipartite graph where citations are directional edges connecting cited and citing publications. For a universe set of total *N* citations, i.e., *N* edges in the bipartite graph, we assume each citation randomly picks one cited publication and one citing publication independently and with equal probability. The random model can also be constructed by randomly permutating the total *N* edges in the bipartite graph. Then specifically for two countries *A* and *B*, the number of citations from *A* to *B* is a random variable denoted as *X*, which follows a hypergeometric distribution *X* ∼ Hyper(*m, n, N*) where parameters *m, n* and *N* have already been explained in Table 1. The probability of observing a value larger (for measuring the over-influence) or smaller (for measuring the under-influence) than *k* can be calculated directly from the hypergeometric distribution, or its approximations, the Binomial distribution or chi-square distribution (Rivals et al., 2006). However, in the context of the citation analysis, the numbers of citations in the contingency table are usually huge and it is very likely to obtain a very tiny *p*-value (e.g., *N* = 139,760,456 in this study). On the other hand, to reject the null hypothesis that two events of *A* citing and *B* being cited are independent is not of interest in this study, while the magnitude of the citation enrichment between the two countries is more relevant. Thus, we only used the enrichment score as the quantitative measurement of the preference of influence between countries while no hypothesis-testing procedure was applied in this study.

### 3.3. Interpreting the enrichment scores

In the previous section, the enrichment score was defined as the logarithm of the citation fold change relative to a random citation process. However, such randomness is conceptual and does not directly correspond to real-world scenarios. In this section, we aim to provide a more intuitive interpretation of the enrichment scores. We first rewrite the enrichment score into the following two forms:

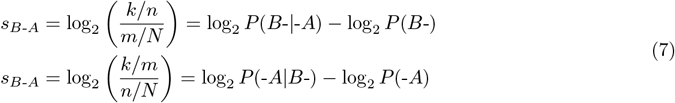

where *B*- and -*A* represent the two events of “*B* being cited” and “*A* citing”, respectively. In Equation 7, for example, *k/n* is interpreted as the conditional probability or the likelihood of *B* being cited given a publication from *A*. In this way, the enrichment score *s*_*B*-*A*_ can be interpreted as the fold increase or decrease of the likelihood of *B* being cited after it is restricted in *A*-citing countries from the globe. Similarly, *s*_*B*-*A*_ can also be interpreted as the fold change of the likelihood of a publication being cited by *A* after the publication is only restricted in *B*-cited countries.

Suppose country *A* cites two countries *B*_1_ and *B*_2_ with corresponding enrichment scores 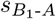 and 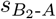. We calculate the difference between the two enrichment scores using Equation 7.

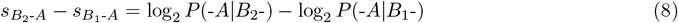

Equation 8 implies, if *A* has a higher enrichment score when it cites *B*_2_ than it cites *B*_1_, given a publication, *A* is more likely to cite it if it is from *B*_2_. In Section 3.1, we demonstrated that mainland China has a higher enrichment score to cite Singapore than the US. Given a publication from Singapore, the conditional probability of mainland China citing it is 0.223, while given a publication from the US, the conditional probability of mainland China citing it is only 0.115. This also shows Singapore has more influence on mainland China on the single-publication level. If both enrichment scores are negative in the comparison, we can state it in the reverse direction: If *A* has a stronger negative enrichment score when it cites *B*_2_ than it cites *B*_1_, *A* is less likely to cite the publication if it is from *B*_2_.

Similarly, if two countries *A*_1_ and *A*_2_ cite the same country *B*, the difference of their enrichment scores is:

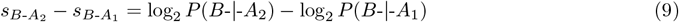

which implies, when *A*_2_ has a higher enrichment score when it cites *B* than *A*_1_ cites *B*, given a publication, *B* is more likely to be cited if the publication is from *A*_2_. Also in Section 3.1, we showed Germany has an over-influence on Austria than the US. Here given the citing country as Austria, Germany is more likely to be cited by it with a likelihood of 0.082, while given the US as the citing country, Germany only has a conditional probability of 0.043 of being cited.

## 4. Results

### 4.1. International citations are dominant

Before performing the citation enrichment analysis, we first analyzed the fractions of country-wise international citations and references. We restricted to countries with a total number of citations and references both larger than 10,000 in the period of this study, which resulted in 82 countries and regions^7^ covering six continents. It is easy to see in Figure 3A, that the majority of the countries have both high fractions of international citations and references. In 79 out of all 82 (96.3%) countries, more than 74% of their citations are international. In 81 out of 82 (98.8%) countries, more than 73% of their references are international. There are only three countries with a small fraction of international citations: 43.9% for mainland China, 48.4% for Ethiopia and 49.9% for the US, thus with a relatively high proportion of domestic citations. Only the US has a small fraction (39.1%) of international references.

**Figure 3.**
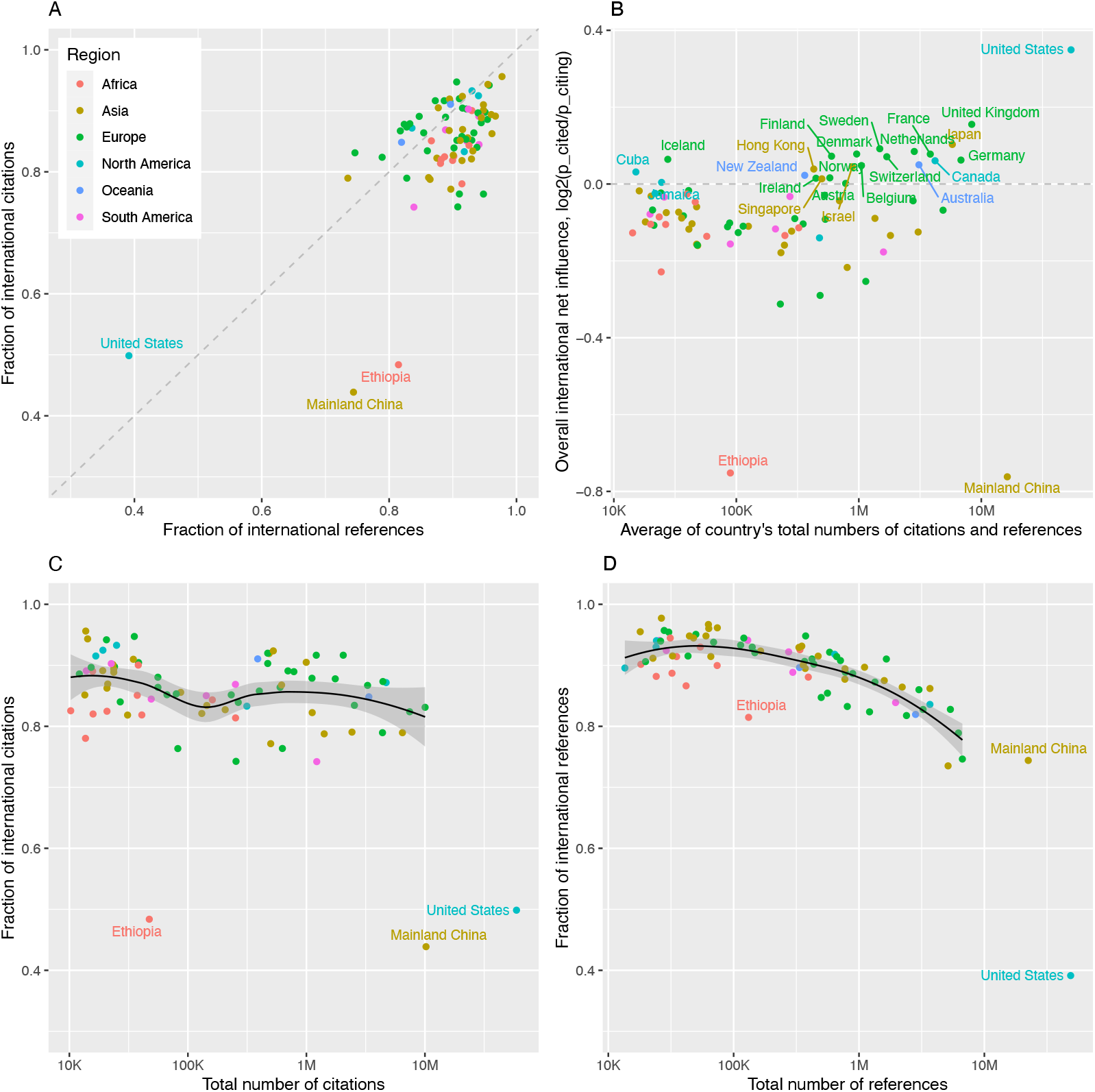
Compare the fractions of international citations and references of 82 countries. A) Scatterplot of the fractions of international references and citations. B) The net international knowledge flows. Countries with values larger than zero are highlighted with their names. C) Relations between the fraction and total number of international citations. D) Relations between the fraction and total number of international references. In Figures C and D, mainland China, Ethiopia and the US were excluded when fitting the loess smoothing lines.

Denote *p*_cited_ and *p*_citing_ as the fractions of international citations and references of a country, we can use log_2_(*p*_cited_*/p*_citing_) to measure the “net” flow of how it receives (the in-flow) or contributes (the out-flow) knowledge to the globe. Figure 3B illustrates that countries with positive net out-flow are mostly highly advanced in science (highlighted with labels), whereas the US has the largest net out-flow internationally. Interestingly, mainland China, as the country with the second-largest number of citations in the globe, has the largest net in-flow, implying its global trend of being significantly influenced by international countries.

Excluding mainland China, Ethiopia and the US which behave differently from other countries, we found the fraction of international citations has a weak negative correlation to the absolute number of total citations of the country (Figure 3C). The fraction of international references also has a negative but stronger correlation to the total number of references (Figure 3D). The decrease in international citations and references may be due to that, a high volume of citations is associated with a high volume of publications which is in turn associated with a high volume of scientific units in the country, i.e., more universities and institutes, which eventually promotes more between-units activities within the country such as collaborations and conferences, and increases the fractions of domestic citations.

As has been described in this section, the fraction-based citation analysis can already reveal important patterns of international influence of countries, however, we argue that such analysis is too general and incomplete. Figure 3B shows the US has the largest net out-knowledge flow internationally. But, we should be careful when concluding that the US is the most significant country that contributes scientific influence to the world. In Section 4.3 where we applied citation enrichment analysis on individual country pairs, we demonstrated that the mutual influences between the US and all other countries in the world are actually under-represented. The net out-flow of the US is due to that the US has a stronger negative preference for citing than being cited by other countries. In other words, the US contributes citations to other countries less than random but it receives citations from other countries even more less. The scenario for mainland China is more complex. With the citation enrichment analysis, we revealed that mainland China’s overall net in-flow is mainly caused by the Western countries where mainland China is cited less than it cites, but the mutual influence between them is under-represented, while it shows over-influence (i.e., more citations than random) partially on science-developing countries, To sum it up, a single aggregated metric on the country-level is not enough for representing the complex patterns of how the country interacts with the globe especially when the interactions are heterogeneous, which will cause insufficient and inaccurate interpretations. Additionally, with the citation enrichment analysis in Section 4.2, we will give a quantitative and theoretical explanation of the negative relations found in Figure 3C-D.

### 4.2. Domestic citations are highly enriched

We calculated the citation enrichment score from one country to the other with a minimum of 100 citations in the period of this study. This resulted in 4,583 country pairs covering 82 countries. Note the country pairs are directional. Figure 4A illustrates the global distribution of the citation enrichment scores. In general, the range of enrichment scores decreases when the number of citations increases. This is a typical pattern of the distribution of log2 fold enrichment where the absolute number of observations follows the hypergeometric distribution (Supplementary File 2).

**Figure 4.**
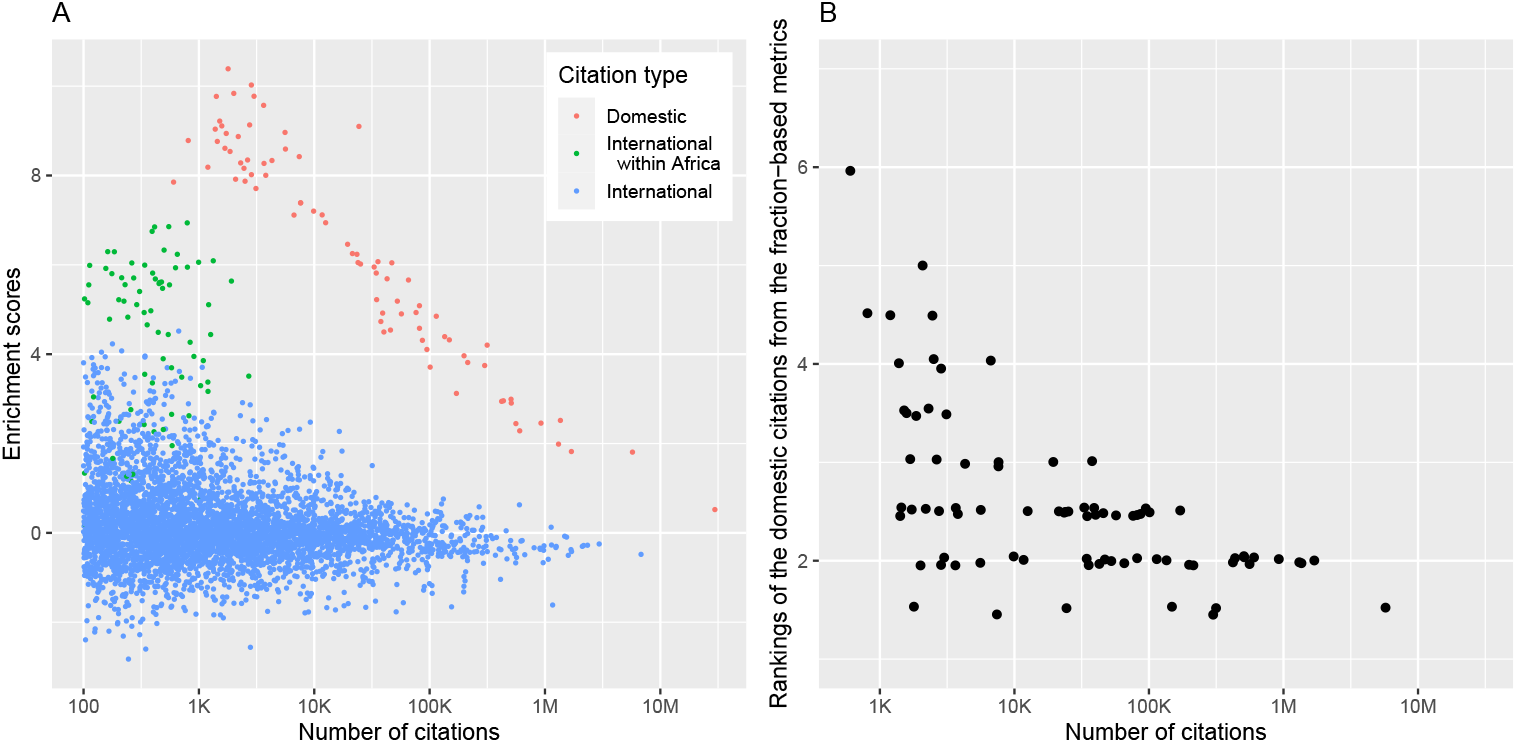
The citation enrichment of all country pairs. A) The scatterplot of the enrichment scores against the numbers of citations. Each dot corresponds to a citation relation from one country to the other. B) Rankings of the domestic citations from the fraction-based metrics. Values on the *y*-axis are the average rankings from the cited countries and citing countries. Each dot corresponds to a country. Small random offsets were added to the *y*-coordinates of points to get rid of overlapping.

We first studied the enrichment of domestic citations, i.e., the preference of of a country citing itself. As shown in Figure 4A, domestic citations are very isolated and show significantly higher enrichment than international citations. Such high enrichment is intuitive to understand because domestic scientific exchanges are expected to be much more active than international ones, which greatly promotes the citations of domestic publications. As described in Section 3.1, if the domestic influence is simply measured by the fraction of domestic citations to the total number of citations or references of that country, as shown in Figure 4B, the ranking of domestic citations drops when the number of domestic citations decreases. In other words, the significance of domestic citations will be much underestimated for small countries. Particularly, the US is the most cited country of all countries, and the most referenced country of 51 countries, and mainland China is the most referenced country of 20 countries. This implies that the US or mainland China will always be evaluated as the most influenced or influencing country if taking the citation fraction as the metric, which is just purely due to their large citation volumes. As an opposite example, the domestic citation of Macau only has an average ranking of 6 by the fraction metric^8^ due to its very small citation volume, but it has a large enrichment score of 7.85 for its domestic citations, representing a very strong preference for citing and being cited by the publications also produced in Macau. Thus, the enrichment method prioritizes the meaningful patterns of scientific influence and may reveal the cognitive patterns in the domestic citation process.

In Figure 4A, it is easy to see that the enrichment score has a well-linear relation to the number of domestic citations in the log scale (the red dots). For domestic citations, we first found that empirically the number of total citations (*m*_*A*-_) or references (*n*_-*A*_) has a linear relation to the number of domestic citations in the log scale (Supplementary File 3), which may be due to the scale-free attribute and the exponential growth of the citation networks (Radicchi and Castellano, 2015). We first fitted the following two linear models on all included countries:

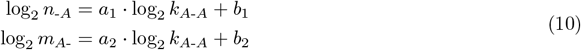

with estimated coefficients *a*_1_ = 0.81, *b*_1_ = 6.20, *a*_2_ = 0.89, and *b*_2_ = 4.45. The *p*-values from the *F* -tests of the two linear regression fits are both significant (*p <* 2.2 × 10^*−*16^, Supplementary File 3).

Let’s denote *T*_*A*-*A*_, *T*_-*A*_ and *T*_*A*-_ as the three citation networks constructed by *A*-domestic, *A*-citing and *A*-cited citations where *T*_-*A*_ and *T*_*A*-_ can be thought of as two background networks of *T*_*A*-*A*_. Since the three networks are assumed to be scale-free, the coefficients of *a*_1_ and *a*_2_ measure the relative growth speed of the networks, i.e. the speed of accepting new citations regardless of any specific countries. The linear fits in Equation 10 can be transformed to 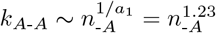 and 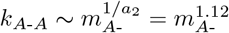. Both power coefficients larger than one implies that *T*_*A*-*A*_ grows faster than its two background networks *T*_-*A*_ and *T*_*A*-_, meaning domestic citation process is more active than in the globe. Also, we can have 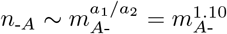, implying the citing network *T*_-*A*_ grows faster than the cited network *T*_*A*-_, thus a faster increasing fraction of domestic references than citations. These results also explain the patterns observed in Figures 3C and 3D.

Then the domestic enrichment score can be written as

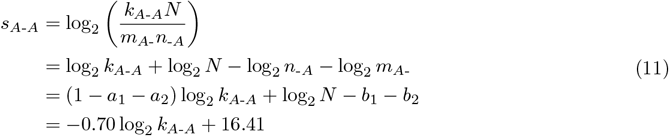

which is the linear regression fit for domestic citations in Figure 4A. Note *N* is the total number of citations of all countries in the analysis, which is fixed and already known.

Next, we explore why the regression coefficient is negative in Equation 11. Let’s simplify the problem to *n*_-*A*_ = *m*_*A*-_, i.e., *A* has the same amount of citations and references for any countries in the analysis, then in the random scenario where two events of “*A* citing itself” and “*A* being cited by itself” are independent, with Equation 5 and 6, we can have the following relations:

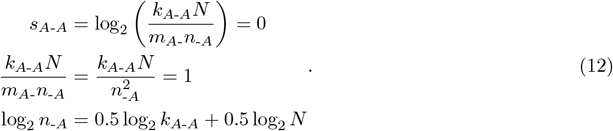

This implies *a*_1_ = 0.5 in Equation 10 and we can obtain *a*_2_ = 0.5 in the same way. The two values of *a*_1_ and *a*_2_ imply that when there is no enrichment, the growth speed of *T*_*A*-*A*_ is quadratic to its background networks, i.e., 1*/a*_1_ = 2 and 1*/a*_2_ = 2, when *N* is fixed. When there is a domestic over-enrichment, more citations from the two background networks are added to *T*_*A*-*A*_, which slows down the growth speed of *T*_*A*-*A*_ to 1*/a*_1_ < 2 and 1*/a*_2_ < 2, and results in *a*_1_ + *a*_2_ *>* 1, which eventually results in the negative trend between the enrichment scores *s*_*A*-*A*_ and log_2_ *k*_*A*-*A*_ (Equation 11). When the domestic enrichment becomes stronger, *a*_1_ + *a*_2_ will be further away from 1, which results in a steeper negative slope for the domestic fit. A complete proof can be found in Supplementary File 3.

With Equation 10, we have

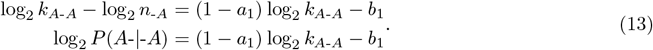

Consider the extreme case where a country only cites itself to reach the maximal enrichment, i.e. *k*_*A*-*A*_ = *m*_*A*-_ = *n*_-*A*_, which yields *a*_1_ = 1. Then the range of the coefficient 1 *− a*_1_ is [0, 0.5) if there is a domestic enrichment. Similarly, we can also have:

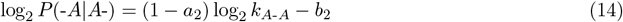

with 1 *− a*_2_ ∈ [0, 0.5). Add these two equations:

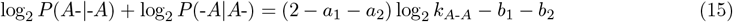

with the coefficient 2 − *a*_1_ −*a*_2_ ∈ [0, 1). This implies, that the likelihood of domestic citations increases as the number of domestic citations increases, but the speed of the increase is slower than the number of domestic citations.

Last, let’s rewrite Equation 12 to:

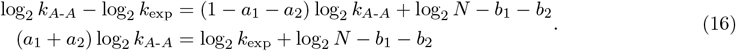

This implies when there is positive domestic enrichment, i.e. *a*_1_ + *a*_2_ *>* 1, *k*_*A*-*A*_ increases slower than its expected version, which results in a decrease of the enrichment scores.

Interestingly, we also found there is a group of green dots between the domestic citations and the main dot cloud in Figure 4A. They are citations, although not domestic, but only between African countries. This implies that the preference of influences between African countries is much stronger than other within-continent international citation relations.

As domestic citations have distinct enrichment patterns from international ones, we excluded them from the analysis in the next sections and we only explored the international citation relations between countries (green box in Figure 1, but excluding the dashed arrows).

### 4.3. Landscape of the preference of international scientific influence

In Section 4.1, we have demonstrated the scientific knowledge flows in the majority of countries are international, which supports the necessity and importance of studying international influence between countries. To construct the global map of pairwise country-level preference of influences, enrichment scores were formatted into a square matrix and visualized by a comprehensive heatmap in Figure 5. In it, rows correspond to the cited countries and columns correspond to the citing countries. Since not every pair of countries has enough citations for calculating the enrichment score, to reduce the sparsity of the matrix, we additionally required each country to have enrichment scores available to at least 24 other countries. Finally, a 72 × 72 square matrix with a sparsity of 22.6% was generated for constructing the world influence map^9^. Note the heatmap is not symmetric, although visually very close to. The mutual influence between two countries, i.e., *s*_*A*-*B*_ and *s*_*B*-*A*_, will be further discussed in Section 4.4.

**Figure 5.**
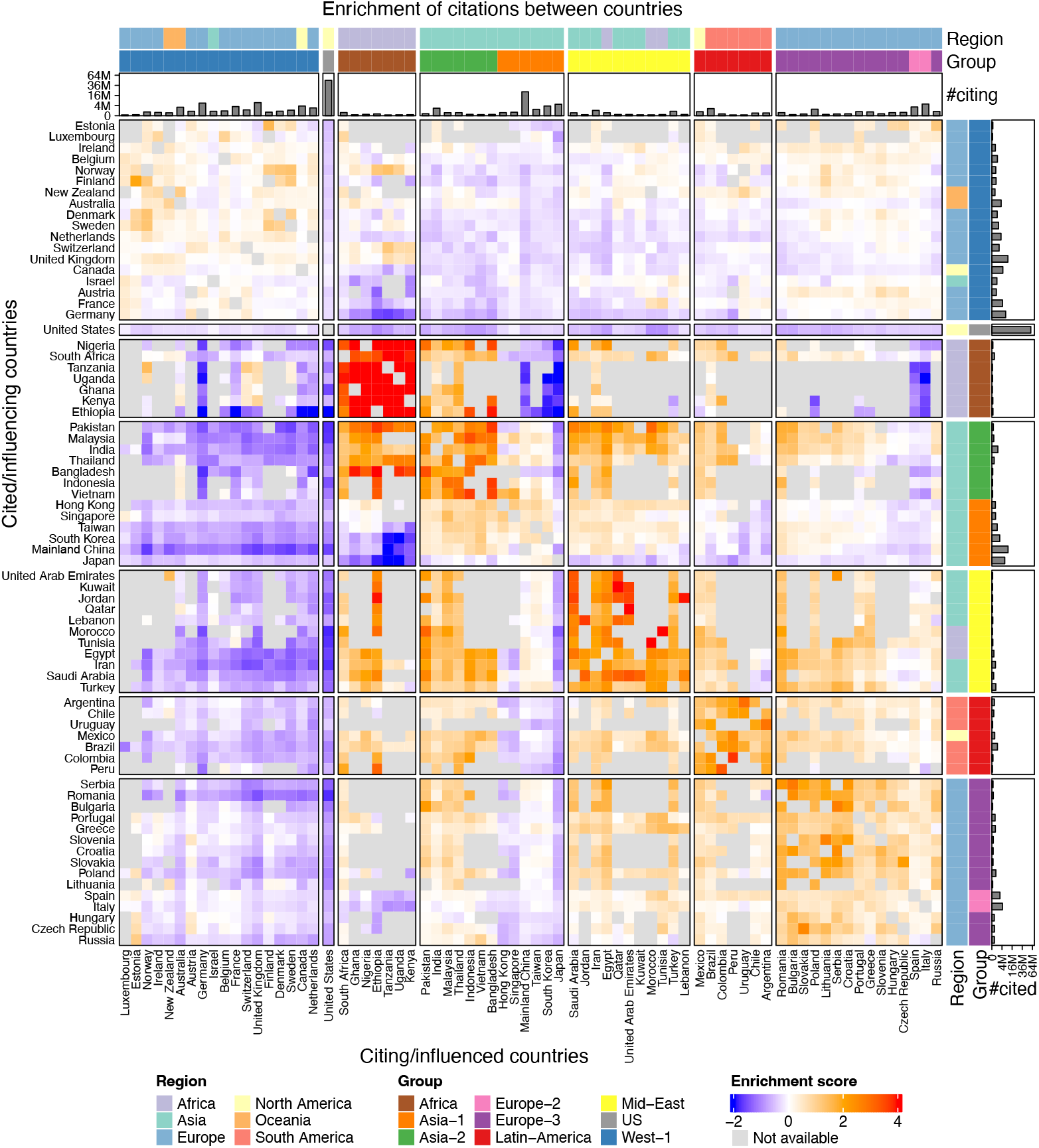
Global map of preference of international scientific influence. Barplots on the two sides of the heatmap are in quadratic scales.

The 72 countries included on the heatmap were partitioned into seven groups both on rows and columns. For the partitioning, we constructed a graph by taking the enrichment matrix as the adjacency matrix and the enrichment scores as the weights of edges. Next, the Louvain graph clustering method (Blondel et al., 2008; Csardi and Nepusz, 2006) was applied to partition the graph. Since the Louvain method does not allow negative values, we only considered country pairs with positive enrichment scores. On the heatmap, rows and columns were further reordered by hierarchical clustering within each group. The separation of the seven country groups can also be validated by the dimension reduction analysis on the influence matrix (Supplementary File 4).

Very surprisingly, the partitioning of 72 countries has a very strong correlation to their geographical classifications or cultural closeness (the “Group” annotation on the heatmap). Countries in the seven groups as well as their geographical classifications are listed in Table 2.

**Table 2:** Country groups identified from the world map of preference of international scientific influence.

| Countries | Geographical classifications | Groups |
| --- | --- | --- |
| Australia, Austria, Belgium, Canada, Denmark, Estonia, Finland, France, Germany, Ireland, Israel, Luxembourg, Netherlands, New Zealand, Norway, Sweden, Switzerland, United Kingdom | North America, Western Europe, Northern Europe, Oceania, Israel | West-1 |
| United States | North America | US |
| Ethiopia, Ghana, Kenya, Nigeria, South Africa, Tanzania, Uganda | Africa | Africa |
| Mainland China, Hong Kong, Japan, Singapore, South Korea, Taiwan | East Asia | Asia-1 |
| Bangladesh, India, Indonesia, Malaysia, Pakistan, Thailand, Vietnam | South Asia, Southeast Asia | Asia-2 |
| Egypt, Iran, Jordan, Kuwait, Lebanon, Morocco, Qatar, Saudi Arabia, Tunisia, Turkey, United Arab Emirates | Middle East | Mid-East |
| Argentina, Brazil, Chile, Colombia, Mexico, Peru, Uruguay | Latin America | Latin-America |
| Italy, Spain | Southern Europe | Europe-2 |
| Bulgaria, Croatia, Czech Republic, Greece, Hungary, Lithuania, Poland, Portugal, Romania, Russia, Serbia, Slovakia, Slovenia | Central Europe, Eastern Europe, Southern Europe | Europe-3 |

In Figure 5 as well as in Table 2, the first two groups contain countries in Northern and Western Europe, developed English countries, and Israel. The common attribute of these countries is that they are highly science-advanced and have closer scientific communications compared to other countries (Leydesdorff et al., 2013; Zimmerman et al., 2009). Countries in both groups can be classified to the so-called highly developed “Western world”. The US shows a slightly overall mutual under-influence on all group-1 countries, thus we only termed group-1 countries as “West-1”, while treating the US as a separated group, emphasizing its uniqueness. However, both West-1 and the US show consistent and strong mutual under-influences on all other countries, implying the scientific recognition between the two parts of the world is weaker than random and the globe is split into two isolated worlds. Although West-1 countries in general show mutual under-influence on the remaining countries on the map, there are still exceptions where individual Western countries influence other countries positively. For example, Switzerland, Norway and the UK have an intermediate mutual over-influence on the three Eastern African countries of Uganda, Kenya and Tanzania (*s* = 0.76)^10^, reflecting their international exchange and cooperation of science (Christine and Falko, 2022). France has an intermediate over-influence on Tunisia and Morocco (*s* = 0.72), which may imply associations between their historical and linguistic relations (Šime, 2023; van Hüllen, 2012).

The third group contains seven African countries. They show the strongest mutual over-influences (*s* = 5.18) compared to the within-group preference of influence in all other groups, implying their strong preference of citing themselves. The high preference of influence within African countries can also be observed in Figure 4A where the green dots are located away from the main dot cloud of international influence. However, the numbers of citations which Africa cites or is cited by other countries in other groups are too small that the majority of the enrichment scores are not available (light grey grids on the heatmap), reflecting their low level of international scientific influence.

The fourth group contains Asian countries. It can be easily observed from the pattern of the preference of influence on the heatmap that they can be further split into two subgroups. The first subgroup contains mainland China, Hong Kong, Japan, Singapore, South Korea and Taiwan, which are more science-advanced with larger citation volumes in the Asian regions, but with relatively lower levels of mutual over-influence (*s* = 0.37). Note these six countries and regions are also culturally similar. Thus, we termed this subgroup as “Asia-1”. Even within the Asia-1 group, we can still see that Hong Kong, Japan and Singapore behave slightly differently from mainland China, South Korea and Taiwan where the former three have weak under-influence from the Middle East and Latin America and they are thought to be closer to Western countries than the latter three ones. Nevertheless, the sub-difference in the Asia-1 group is small compared to the global trend, thus Asia-1 was not further split on the map. The second subgroup contains countries in South Asia and Southeast Asia which are less science-advanced in the regions but show higher levels of mutual over-influence (*s* = 2.48). We termed this subgroup as “Asia-2”.

The fifth group includes countries from the Middle East or the pan-Arab world. The sixth group includes countries from Latin America. They show consistent mutual over-influence within groups (*s* = 2.34 and 1.80).

The last group contains countries from Southern, Central and Eastern Europe. They are also manually split into two subgroups based on the patterns on the heatmap. We termed Spain and Italy as “Europe-2” and other group-7 countries as “Europe-3”. It might be unexpected that Spain and Italy are thought to be highly developed European countries, but under the context of citation enrichment, they indeed show an under-influence on West-1 (*s* = *−* 0.17) and an over-influence on the Middle East and Europe-3 (*s* = 0.33 and 0.51), making them separated from the Western world. This result reveals the imbalance of scientific influence within the European Union. The heatmap also reveals Spain has a stronger mutual over-influence on Latin America than Italy (*s* = 0.56 vs. 0.06), which may relate to the historical relations between Spain and Latin America. Interestingly, we found Spain has a slightly stronger over-influence on Spanish-speaking Latin American countries (*s* = 0.59) than on Brazil (Portuguese-speaking, *s* = 0.37), while a higher over-influence was found for Portugal on Brazil (*s* = 0.86). These results may reflect the strong associations between scientific influences and cultural and linguistic similarity (Echeverría-King et al., 2023; Narváez-Berthelemot et al., 1993; Almeida and Correia, 2020).

The Louvain graph clustering method expects an undirected graph, while the enrichment matrix contains directional relations. To aggregate two enrichment scores *s*_*A*-*B*_ and *s*_*B*-*A*_ for a pair of countries, three methods can be used: taking the sum, the maximum or the minimum of the two values. In the clustering step, there is also a pre-selected parameter named “resolution” which controls the tightness of the clusters. To test the robustness of the partitioning, we enumerated all combinations of the aggregation methods and the resolution parameter. As shown in Supplementary File 4, the partitioning of the seven groups is almost stable among different combinations of parameters. We found Bangladesh and Pakistan are classified into the Africa group in some clustering iterations, which may imply their relatively strong scientific relations to Africa in research fields like infectious diseases, agriculture, food, and public health (Bidaisee and Macpherson, 2014). Similarly, Morocco and Tunisia are partially classified into an eighth group, suggesting associations with their identities as Arabic countries but with comparatively weaker connections to the Middle East (Colombo et al., 2018).

#### 4.3.1. Between-group influence: two separated worlds

To better summarize the world map of the preference of scientific influence, we constructed a simplified group-level influence network. The preference of influence between the two groups was calculated as the average enrichment scores of their member countries. To highlight both strong over- and under-influences, we only considered group pairs with absolute enrichment scores larger than 0.15. To reduce the complexity of interpreting the influence network, we separated it into an under-influence network and an over-influence network, as shown in Figures 6A and 6B.

**Figure 6.**
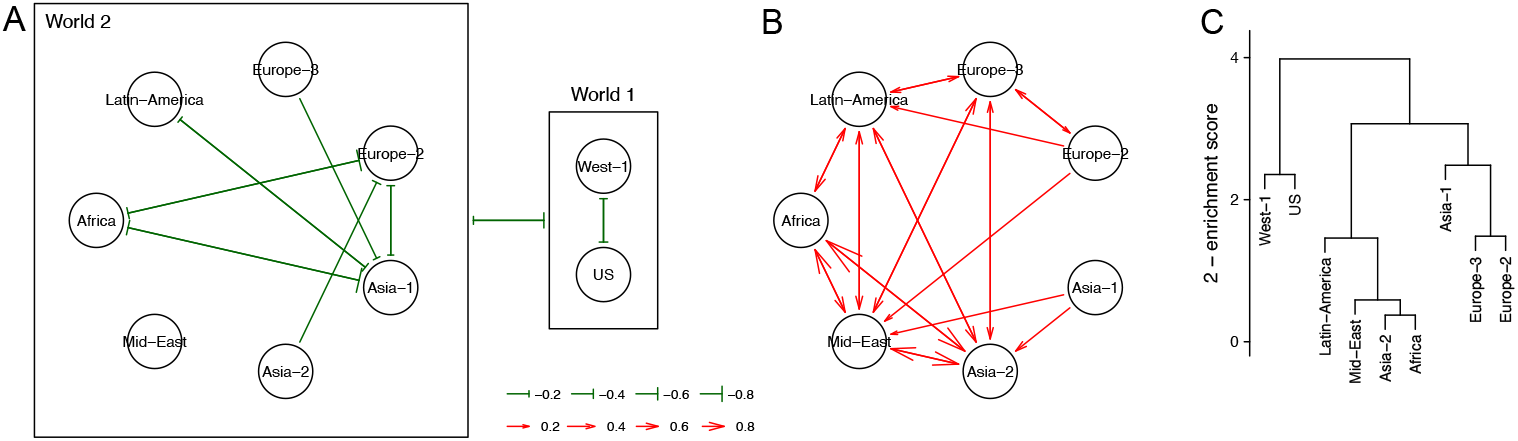
Simplified network of the preference of scientific influence. A) The under-influence network. B) The over-influence network. C) Hierarchical relations of country groups based on their pairwise preference of influences. A subtree with a height less than 2 can be interpreted as the average mutual over-influence of its member country groups.

The nine country groups can be put into two broad categories which we termed as “World-1” and “World-2”. The World-1 category contains West-1 and the US, which show consistent mutual and strong under-influences on the countries in World-2 (*s* = − 0.74 for World-2 on World-1 and *s* = − 0.30 for World-1 on World-2, Figure 6A). In World-1, the US shows an intermediate level of mutual under-influence on West-1 countries (*s* = − 0.35 for West-1 on the US and *s* = − 0.28 for the US on West-1), but it is mainly due to the high domestic citations in the US.

The remaining countries compose World-2 and they are quite separated from World-1. In general, countries in World-2 have more mutual over-influence. Especially, Africa, Asia-2 and Mid-East show the strongest mutual over-influences (*s* = 2.20, Figure 6B), implying their strong preference of citing publications in their regions. There are still regional under-influences sourced from Europe-2 and Asia-1. Interestingly, Europe-2 and Asia-1 show certain levels of exclusive enrichment patterns in World-2: Europe-2 has a mutual over-influence on Europe-3 (*s* = 0.40) and contributes an over-influence on Latin-America (*s* = − 0.31) while Asia-1 has under-influences on both (*s* = − 0.22 and −0.26); Asia-1 has an over-influence on Asia-2 (*s* = 0.56) while Europe-2 receives an under-influence from it (*s* = −0.39). The mutual influence between Europe-2 and Asia-1 themselves is *s* = − 0.24. This pattern may imply Europe-2 and Asia-1 are local influencers in their corresponding regions. However, Europe-2 and Asia-1 show over-influences both on Mid-East (*s* = 0.33 and 0.23) and mutual under-influence both on Africa (*s* = − 0.86 and − 1.09).

Specifically for West-1, Europe-2 and Europe-3, West-1 exhibits an intermediate under-influence on Europe-2 (*s* = *−* 0.17) and a stronger under-influence on Europe-3 (*s* = *−* 0.37) while it receives almost neutral influence from both Europe-2 (*s* = 0.057) and Europe-3 (*s* = 0.089). This suggests an imbalance, delay and isolation in the scientific knowledge flow within the Western world and Europe.

It is interesting to see, Europe-2 and Asia-1, which are the two most science-advanced groups in World-2, contribute partial under-influences to other countries in the regions. Together with the separation of World-1 from World-2, we could infer that the dissemination of scientific knowledge is faster between science-advanced countries if they are tightly related, while slower to reach science-developing countries, which eventually causes the separation between them on the world influence map.

Figure 6C illustrates the hierarchical relations of the nine country groups based on their average citation enrichment. The dendrogram confirms the separation of World-1 and World-2, the clustering of Europe-2 and Europe-3, and the clustering of Latin-America, Mid-East, Asia-2 and Africa.

#### 4.3.2. Within-group influence: small worlds

Within individual country groups, we also observed “small-world” patterns, where countries inside can be further split into subgroups. As already demonstrated in Figure 5, by directly observing the influence patterns on the heatmap, Asian countries are split into Asia-1 and Asia-2, and Southern, Central and Eastern European countries are split into Europe-2 and Europe-3.

West-1 shows a certain level of heterogeneity of mutual enrichment. Similar to the partitioning analysis of the global map, we applied the Louvain method only on the 18 West-1 countries, generating three subgroups both on rows and columns (Figure 7A). We found that countries also exhibit geographical or cultural similarities within subgroups. The first subgroup contains the five Northern European countries which show stronger mutual over-influence (*s* = 1.05) than other West-1 countries, particularly between Estonia and Finland (*s* = 1.76). The second subgroup contains five English countries (*s* = 0.40), particularly with high mutual over-influence between Australia and New Zealand (*s* = 1.04). The third subgroup mainly contains mainland European countries (*s* = 0.29, excluding Israel). Northern Europe has a consistent mutual over-influence on English countries (*s* = 0.19). Interestingly, in mainland Europe, Netherlands, Luxembourg and Belgium show relatively stronger mutual over-influence on northern Europe (*s* = 0.26) and English countries (*s* = 0.12), as their geographical locations as bridges connecting the regions. However, as a comparison, the other four mainland European countries, i.e., France, Germany, Switzerland and Austria, show almost-neutral mutual influence on English countries (*s* = *−* 0.07) and Northern Europe (*s* = 0.06), implying the four countries are less preferable to cite or be cited by these two regions. Last, it can be observed that being different from other Northern European countries, Estonia also receives over-influence from mainland Europe (*s* = 0.23), implying its role as an active knowledge receiver as it is also located on the European mainland. Nevertheless, such a small-world pattern of countries is mainly observable within West-1. The country-subgroups can not be well distinguished from how they interact with World-2 countries (Supplementary File 5).

**Figure 7.**
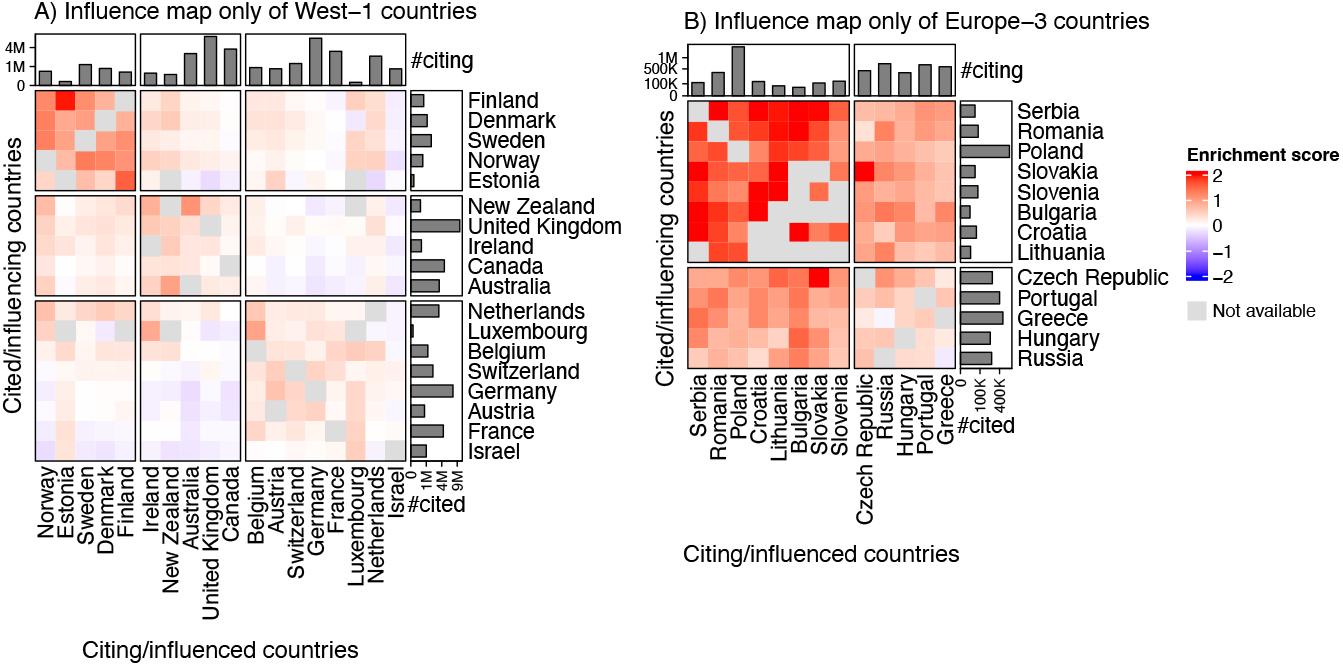
Preference of international influcence of A) 18 West-1 countries and B) 13 Europe-3 countries. Barplots on the two sides of the heatmap are in quadratic scales.

Although countries in Europe-3 show overall mutual over-influences, a certain level of heterogeneity can still be observed. In Figure 7B, since almost all mutual influences are positive, making the graph clustering approach less effective, we partitioned countries simply by hierarchical clustering. We found a core set of 8 countries (Serbia, Romania, Poland, Slovakia, Slovenia, Bulgaria, Croatia and Lithuania) with stronger mutual over-influence (*s* = 1.80 vs. 0.87 for the rest) and they have consistently smaller citation volumes (except Poland) than other countries in Europe-3. Interestingly, these eight countries share certain cultural and historical similarities, as all are Slavic countries, which may also imply the association with their science dissemination patterns. Finally, it is worth noting that the Czech Republic and Slovakia exhibit a strong mutual over-influence (*s* = 2.36), likely associated with their historical union.

### 4.4. Balance of mutual citation enrichment

The country-to-country influence is directional. A country contributes influences to other countries and at the same time it receives influences from them. Specifically for two countries *A* and *B*, the mutual preference of influences are *s*_*A*-*B*_ and *s*_*B*-*A*_. If *A* has the same level of preference of influencing *B* as it is influenced, measured by *s*_*A*-*B*_ = *s*_*B*-*A*_, we could say that the mutual preference of influence between *A* and *B* is balanced. As we have shown countries in the same country group share very similar enrichment patterns, also to reduce the complexity of the interpretation, in this section, we studied the preference of influence balance in each country group only by selecting a representative country. Additionally, by observing the balance patterns, Asia-1 is additionally split into Asia-1-1 (mainland China, South Korea, and Taiwan) and Asia-1-2 (Hong Kong, Japan, and Singapore). Africa was removed from the analysis because the enrichment scores are missing between it and most of the other countries. We selected the following nine countries: the US, Netherlands (West-1), Spain (Europe-2), Poland (Europe-3), Brazil (Latin-America), Saudi Arabia (Mid-East), Thailand (Asia-3), mainland China (Asia-1-1) and Hong Kong (Asia-1-2). Figure 8 illustrates the mutual influences of the selected countries on the globe. The complete analysis for all individual countries can be found in Supplementary File 6.

**Figure 8.**
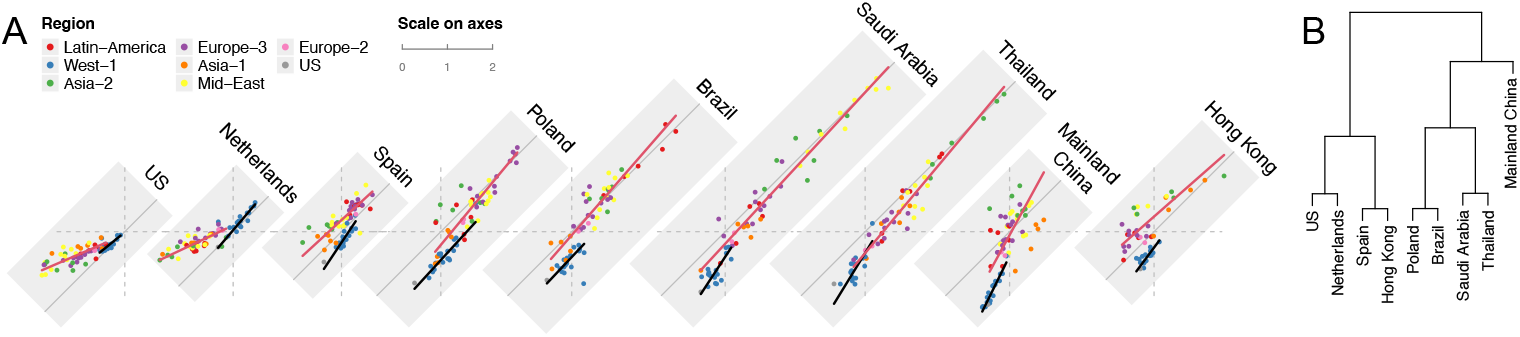
Preference of influence balance of the nine selected countries. A) Each grey rectangle corresponds to the plotting panel of a country. *s*_*B*-*A*_ corresponds to values on the *x*-axis and *s*_*A*-*B*_ corresponds to values on the *y*-axis where *A* corresponds to one of the nine selected countries and *B* corresponds to individual countries in each plotting panel. All nine panels are aligned by their *x*-axes and have parallel *y*-axes. The scales on the *x*-axes and *y*-axes are the same for all panels. The linear fit is applied to World-1 and World-2 separately. B) Hierarchical clustering on the citation balance patterns in the two worlds. Africa was removed from the analysis.

In Figure 8, most data points are located close to the line *y* = *x*, which implies that in general the mutual preference of influence is balanced for these countries. This pattern can also be observed from Figure 5 where the lower triangle shows similar patterns as the upper triangle on the square heatmap. We found, that for each country in Figure 8, the patterns of mutual preference of influence are approximately linear over countries, and the linear relations are distinct between World-1 and World-2. In the citation balance analysis, we took *y* = *x* (completely balanced) as the baseline, thus, we first fitted the ordinary linear model *y* −*x* ∼ *y* + *x*, which is identical to minimizing the sum of the squared distances of data points to *y* = *x* in the fit, then the fitted equation was transformed to the form of *y* = *αx* + *β*.

To better explain the slope coefficient *α*, let’s take country *A* as a fixed country and *B* ∈ *W* where *W* is the set of countries in World-1 or World-2. The linear model between *s*_*A*-*B*_ (the enrichment score of *A* being cited) and *s*_*B*-*A*_ (the enrichment score of *A* citing) is

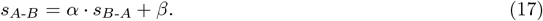

Simply speaking, *α* corresponds to the increasing rate of *s*_*A*-*B*_ against *s*_*B*-*A*_. When *α >* 1, *s*_*A*-*B*_ increases faster than *s*_*B*-*A*_. Let’s assume there are two countries *B*_1_ and *B*_2_ with 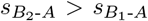, i.e., *A* receives a stronger citation enrichment from *B*_2_ than *B*_1_. With Equations 5, 6 and 17, there are

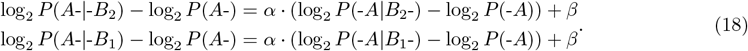

Subtracting Equations 18 on both sides yields:

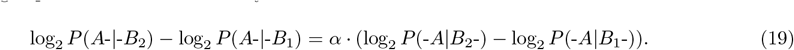

The left side of Equation 19 is the changing rate of the likelihood of *A* being cited if the given citing country is changed from *B*_1_ to *B*_2_. Similarly, the right side of Equation 19 corresponds to the changing rate of the likelihood of *A* citing given the cited country is changed from *B*_1_ to *B*_2_. Let’ denote:

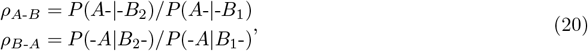

then Equation 19 is simplified to:

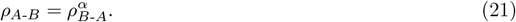

With the condition 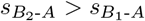, we can have *ρ*_*B*-*A*_ > 1. According to Figure 8, the slope coefficients for all linear regressions are positive, then we can also have *ρ*_*A*-*B*_ *>* 1. Then with Equation 21 we can have the following relations:

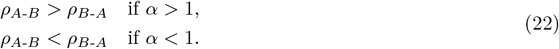

Equation 22 implies, when the enrichment score of *A* citing the world *W* increases, if *α* > 1, the likelihood of *A* being cited by *W* increases faster than *A* citing *W*, or we could say *A* expands its influence faster to *W* than receiving from it. Similarly, if *α* < 1, we could say that the likelihood of *A* citing *W* increases faster than *A* being cited by *W*, or we could say *A* receives the influence from *W* faster than expanding to it.

Particularly, when the enrichment scores are negative, the decrease of the enrichment scores implies the strengthening of the under-influence. Let 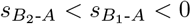, then we can have *ρ*_*A*-*B*_ < 1 and *ρ*_*B*-*A*_ < 1, and the following relations:

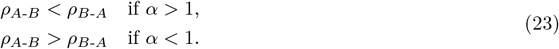

This implies, that when the under-influence becomes stronger from the world *W*, if *α >* 1, the likelihood of *A* citing *W* decreases faster than *A* being cited by *W*, or we could say *W* has a faster loss of its influence on *A*. If *α <* 1, the likelihood of *A* being cited by *W* decreases faster than *A* citing *W*, or we could say *A* has a faster loss of its influence on *W*.

The intercept coefficient *β* measures the citation enrichment of *A* being cited by *B* when *A* neutrally cites *B*, i.e. *β* = *s*_*A*-*B*_, when *s*_*B*-*A*_ = 0. It can be inferred as the systematic imbalance of the mutual preference of influences between *A* and *W*.

The overall preference of influence balance of *A* on *W* denoted as *η*_*A*_ is simply defined as:

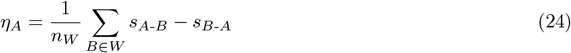

where *n*_*W*_ is the number of countries in *W*. A positive value of *η*_*A*_ represents the net preference of *A* contributing influence on *W* (let’s call it the net out-influence for simplicity). Note when *α* ≈ 1, then *η* ≈ *β*, i.e., the systematic imbalance.

The coefficients of the linear fits as well as the overall influence balance for the nine countries on World-1 and World-2 are listed in Table 3. In Figure 3B, we have demonstrated the overall trend of the US is exporting its scientific knowledge to the world. Here with the citation balance analysis, more detailed internal structures can be revealed. First, in Figure 8 it is straightforward to see, that the net out-influence of the US is mainly from World-2 (*η* = 0.56) where corresponding countries are all above *y* = *x*, while it is almost balanced on West-1 (*α* = 0.82, *η* = 0.07). Second, notice all the enrichment scores between the US and World-2 are negative, this means the US citing World-2 is even more under-represented than World-2 citing the US, which results in the net out-influence from the US. Thus, a more proper interpretation on “the net out-influence from the US” is not “the US exports knowledge to World-2”, rather “the knowledge dissemination is more restricted from World-2 to the US”. Third, the slope coefficient is far less than 1 (*α* = 0.43) on World-2. According to Equation 23, it can be interpreted as when the under-influence from the US gets stronger on World-2, the US has a faster loss of its influence, i.e. a faster decrease of the likelihood of World-2 citing the US. Netherlands has similar patterns to the US of interacting World-2.

**Table 3:**
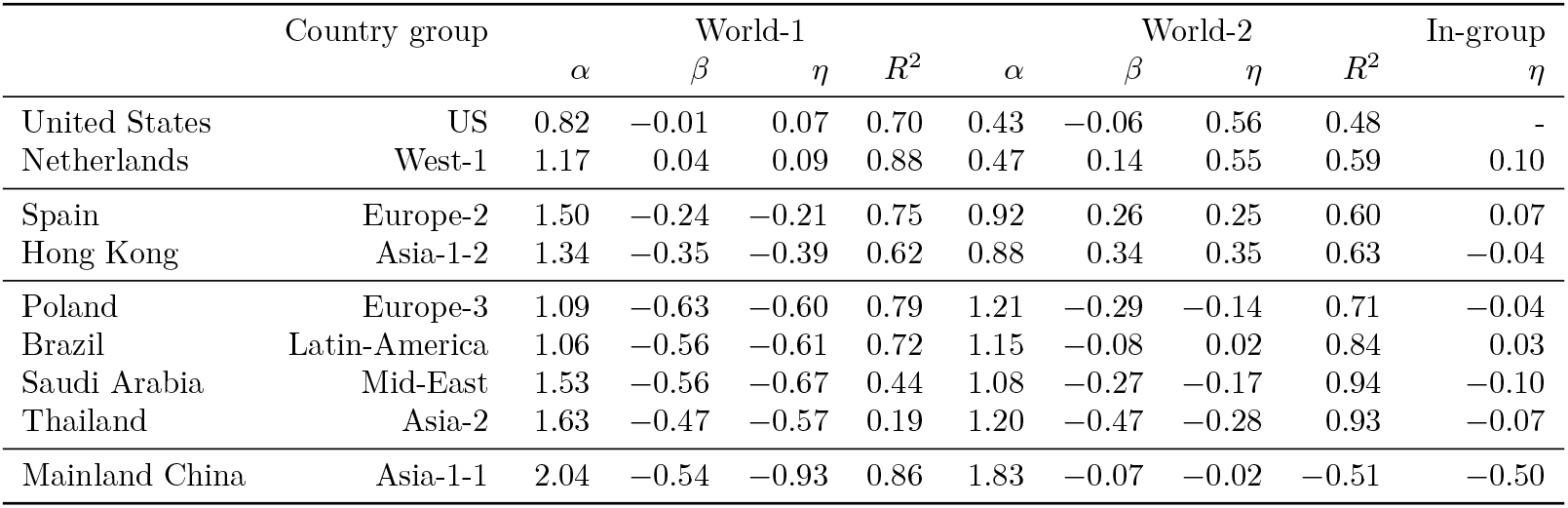
Coefficients of the linear fits as well as the overall influence balance in the citation balance analysis. *α*: slope coefficient, *β*: intercept coefficient, *η*: overall influence balance, *R*^2^: R-squared statistic from the linear regression. The *R*^2^ of the linear regression of mainland China on World-2 is negative because the fitting is not directly from ordinary linear regression, and it also implies a highly unreliable fit. The last column: the overall influence balance of the country to other countries within the same group. The splitting of countries is based on the clustering in Figure 8B.

Mainland China is another extreme country in Figure 3B which receives the largest in-knowledge flow from the world. Here as revealed in Figure 8, the citation influence balance pattern for mainland China is more complex, where it has a stable net in-influence flow from World-1 (*η* = − 0.93) but a heterogeneous and partial net out-influence on World-2. On World-1, mainland China has the largest slope coefficient (*α* = 2.04) in the nine countries, with all the enrichment scores being negative. We conclude the likelihood of mainland China citing World-1 decreases faster than cited, or we could say World-1 loses its influence on mainland China faster if the under-influence between them becomes stronger. The intercept coefficient *β* = − 0.54 implies a systematic offset where even when mainland China neutrally cites World-1 with zero enrichment, there is still a strong under-representation of mainland China’s influence on World-1. The balance between mainland China and World-2 is very heterogeneous and there is no reliable linear relation of the mutual influence if taking World-2 as a whole (with a negative *R*^2^ of − 0.51^11^). However, if inspecting individual country groups, we can still observe that similar to World-1, mainland China has a net in-influence and a systematic under-influence on Asia-1 (*η* = − 0.67, *β* = −1.18, Supplementary File 7) with a slope coefficient *α* = 1.99. As Asia-1 is also classified as a science-advanced region in Asia, the similar coefficients from the balance analysis may imply similar mechanisms of the mutual citation processes between mainland China and Asia-1/World-1. Nevertheless, in general, World-1 has an under-influence on mainland China while Asia-1 has an over-influence on it, indicating a system-level preference of influence in the two regions on mainland China. For other country groups, mainland China has almost balanced mutual citations with Mid-East (*η* = 0.02, *β* = − 0.12) and Europe-3 (*η* = 0.04, *β* = 0.19), but both have large slop coefficients (*α* = 1.93 and 2.96 respectively, Supplementary File 7), implying mainland China expanding its influence faster to these two regions as the mutual over-influence increases.

Interestingly, we found that, although Spain and Hong Kong are categorized into World-2, under the context of influence balance analysis, they show reverse mutual influence patterns to other World-2 countries. When they interact World-2, we found they have net out-influence (*η* = 0.25 and 0.35) while other selected World-2 countries have close-to-zero or overall in-influence to World-2. It implies that Spain and Hong Kong are active knowledge contributors in World-2. Spain and Hong Kong also have positive intercept coefficients (*β* = 0.26 and 0.34) on World-2, reflecting the positive preference of influencing World-2 on the baseline level. As a comparison, when interacting with World-1, we found it is a common pattern that all selected World-2 countries have net in-influence from World-1 (the values of *η* are all negative) and mutually exclusive influence on the system-level (the values of *β* are all negative). However, Spain and Hong Kong have weaker imbalance and exclusivity from World-1 than other World-2 countries (*β* = − 0.24 and − 0.35 vs. − 0.55, *η* = − 0.21 and − 0.39 vs. − 0.68)^12^.

We also investigated the overall influence balance of a country within its own group. For example, the average preference of mutual influence balance of Netherlands on all other 17 West-1 countries is 0.10. As shown in the last column of Table 3, the preference of mutual scientific influence is almost balanced inside all country groups, except Asia-1-1 which is mainly because mainland China has a huge imbalance of citing and being cited by international publications (Figure 3A-B). However, South Korea and Taiwan which are also in Asia-1-1, are still very mutually balanced (*η* = 0.02).

Figure 8B illustrates the relations of the nine countries by hierarchical clustering on the regression coefficients from the linear fits on World-1 and World-2. Compared to the dendrogram in Figure 6C which is based on the static world influence patterns, in the citation balance analysis which additionally considers the dynamical relation of a country to a world, we found Poland, Brazil, Saudi Arabia and Thailand, which represent Europe-3, Latin-America, Mid-East and Asia-2, are still tightly clustered, but, Spain and Hong Kong, which represent partial Europe-2 and Asia-1, are now clustered together and closer to World-1. This implies, that although statically, Spain and Hong Kong fall within World-2 due to their relatively lower fold citation enrichment to World-1, their dynamics of interacting with the globe are closer to World-1. The separation of Spain and Hong Kong from World-1 is more due to a systematic imbalance.

## 5. Discussion and conclusion

The citation enrichment analysis presented in this study, being different from traditional bibliometrics methods, provides a new aspect of evaluating scientific influence among countries. We demonstrated the advantages of citation enrichment analysis in that it reveals meaningful and well-structured patterns of global scientific influences that are missing or have not been well-explored in current studies. The citation enrichment metric is built on a well-defined statistical framework and has a clear meaning for interpretation. We believe the citation enrichment analysis will be a powerful tool for bibliometrics studies to advance our understanding of global knowledge dissemination dynamics.

Similar forms of citation enrichment have been mentioned in literature. On the publication level, Balland and Boschma (2022) and Fontana et al. (2019) applied a metric named “Relative Scientific Advantage” to measure whether publications from a country are enriched in a specific research topic, i.e., to evaluate country-topic associations. A positive enrichment is called a specialization of the country on the research topic. Finardi and Buratti (2016) applied a metric named “Probability Affinity Index” to study the levels of collaboration between two countries. Nevertheless, the framework of the enrichment analysis proposed in this study is more general and extensible, and it can be applied to a wide range of research topics.

### 5.1. Implications

The citation enrichment metric provides a complementary view of global science influence. It has several advantages for bibliometrics studies. First, it is a relative metric which evaluates influence on a normalized level, making it suitable for studying and comparing both large and small science countries under the same scale. Second, by comparing actual citation patterns with a statistical random citation model, the metric may capture researchers’ cognitive processes when selecting citations. Third, the citation enrichment metric is directional, which largely extends the scope of analysis. It allows researchers to explore not only which countries are influential but also how their influence is distributed and received, uncovering complex interconnections and patterns of global knowledge exchange.

Our empirical analysis has revealed varying degrees of isolation in global scientific influence. Such isolation may reflect different levels of delay in science dissemination, especially from science-advanced regions to science-developing regions. We observed a significant enrichment of domestic citations, indicating that scientific knowledge tends to disseminate more quickly within countries. This could be attributed to the accessibility of local research, cultural alignment, and shared institutional frameworks that prioritize national over international scientific influence. Internationally, there is an obvious exclusivity of mutual preference of influence between science-advanced and developing countries. This may imply that science is more restricted within science-advanced countries and it has a much faster and stronger dissemination than to the other parts of the globe. Specifically, in the clustering analysis, we found that preferences in scientific influence align strongly with geographical and cultural proximity. Such associations were observed on a large scale, such as in Europe, the Middle East or Latin America, reflecting their preference of regional scientific exchange. The citation enrichment analysis also revealed such associations for countries geographically distal but in close historical, linguistic, or cultural relations, such as France with Morocco/Tunisia, or Spain with Spanish-speaking Latin-American countries. These results highlight the sensitivity of the citation enrichment metric. The associations on various levels provide strong evidence for further studies on the interplay between geographical proximity, cultural connections, and scientific exchange.

### 5.2. Limitations

The citation enrichment metric only utilizes the total numbers of citations of countries in the model, thus, it has the common limitations as other citation-based bibliometrics methods. Specifically, it does not account for the diverse citation patterns that arise due to variations across research fields, journals, institutions, or the impact of individual publications (Waltman and van Eck, 2013). This may limit a comprehensive understanding of the complex patterns in the citation process. Scientific influence often varies by discipline. When analyzing the citation dynamics between two countries, it is critical to ask which research fields show the strongest mutual citation preference. This question is particularly significant for countries with smaller science volumes, which may specialize in specific disciplines rather than having a broad research spectrum. Citation behaviours may also differ between high-impact and lower-impact journals. For example, one might investigate whether mutual citation preferences are stronger in prestigious journals or in more specialized publications.

The empirical analysis presented in this study is based on a static snapshot of the PubMed data. However, our study makes a hypothetical inference on citation dynamics, suggesting that scientific knowledge tends to reach domestic researchers and closely related countries more quickly, while with a delay in dissemination to other regions. In the citation balance analysis, countries grouped together exhibit shared attributes in the expansion of their citation networks. This observation was interpreted as reflecting different stages of network growth, where countries with larger citation volumes may represent more mature stages of network expansion. Nevertheless, the current analysis in this study does not directly capture the dynamics of citation networks over time. Thus, longitudinal studies are essentially required, which could provide direct evidence of how scientific influence grows and spreads among countries.

Scientific collaboration is an important topic in bibliometrics studies (Sonnenwald, 2007), which represents a more direct type of influence of which researchers work together to produce new knowledge. In this work, we only focused on publications uniquely associated with a single country (domestic studies) while excluding publications from international collaborations (the grey rectangle in Figure 1). Nevertheless, bibliometrics studies have revealed positive correlations between collaborations and mutual citations (Milojević, 2010; Liu et al., 2023; Aman, 2016), as collaborations promote the visibility of co-authors’ research to wider communities, and strengthen the preference of citing their works. Thus, a combination of citation and collaboration analysis can help to explore how collaborative efforts directly influence citation preferences and their dynamic relationships.

In Section 5.4, we will discuss how such proposed analyses can be applied under the general framework of citation enrichment analysis.

### 5.3. Methodology

The citation enrichment metric is defined on citations, not on publications, which should be noted when performing interpretations. Under the random citation process, each citation has the same probability of being picked. Then, on the publication level, the probability of picking a publication is proportional to the citations it receives. Citation networks are normally modelled as scale-free networks, where the node degree distribution follows the power-law distribution. One of the standard random models to generate scale-free networks, the Barabasi-Albert model (Albert and Barabási, 2002), expands the network in a way that the probability of a node (publication) accepting new links (citations) is proportional to its degree (number of citations it has already received). Thus, the random citation model preserves these structural properties of the citation network.

The citation enrichment metric proposed in this study simply divides the observed number of citations by the expected one without considering the variance if treating the number of citations as a random variable *X*. There is another widely used method in statistics that measures the relative distance of an observed value of *X* to its null distribution, the *z*-score, defined as *z* = (*k − k*_exp_)*/σ* where *σ* is the standard deviation of *X* from the hypergeometric distribution. In Supplementary File 8, we demonstrated that using *z*-scores generates very similar country classifications as in Section 4.3, but the variance of the *z*-scores now has a positive dependency on the number of citations. Nevertheless, the form of fold enrichment has a clear meaning for interpretation and is more convenient for further statistical analysis.

The universe set *U* is also an important component of the framework of citation enrichment analysis from two aspects. First, all categories in the 2 × 2 contingency table are restricted within the universe set. Second, the enrichment score is a relative metric against a random process which is performed within the universe. The universe defines the context of *what are the citations enriched against*. It should have a clear meaning and the interpretation of the enrichment should not be made beyond the scope of the universe. In this study, the complete statement on the enrichment of *A* citing *B* should be “*A* has an over-influence from *B over the global average*”. In the empirical analysis we performed, we observed that the US has an overall under-influence even on closely related West-1 countries, due to the high volume of its domestic citations. To get rid of such an effect, we can redefine *U* to include only international citations (excluding domestic ones). As shown in Supplementary File 9, the US now has mutual over-influences on West-1, but we should explicitly state that such enrichment is over the international average. Choosing different *U* allows for the exploration of citation behaviours across various levels. For example, we can study the citation process within Europe by restricting *U* to European citations.

### 5.4. Extension

From the aspect of methodology, the enrichment score measures the association between two events represented in the row and column of the 2 × 2 contingency table, restricted in a certain universe set. By choosing different events and universes, the enrichment analysis can be generalized as a powerful tool for bibliometrics analysis. Let’s denote the row and column events as *E*_*a*_ and *E*_*b*_ respectively, and the universe set as *U*. In this study, *E*_*a*_ is “country *A* cites”, *E*_*b*_ is “country *B* is cited”, *U* is the set of total citations in the globe. As example, we can choose other events for *E*_*b*_ to study international influences on various levels:

- Journal-to-country: *E*_*b*_ is “a journal is cited”.
- Institution-to-country: *E*_*b*_ is “an institution is cited”.
- Author-to-country: *E*_*b*_ is “an author is cited”.
- Publication-to-country: *E*_*b*_ is “a publication is cited”.

Then specifically, we can answer the question: Does a journal/institution/author/publication have general international influence or is its influence only restricted to one or a few countries? For small units such as authors or publications, a cutoff of the minimal number of citations might need to be set to ensure the interpretation of the enrichment score is reliable.

Under the same framework, the citation enrichment metric can also be extended to longitudinal studies, e.g., in the following two directions:

- Define *E*_*a*_ as “country *A* cites”, *E*_*b*_ as “country *B* is cited only for its publications published on year *y*”, and *U* as the set of total citations on and after year *y*. By applying the citation enrichment analysis to a list of continuous years, we can study the longitudinal trend of country-to-country influences, and answer the question “are the two worlds becoming more separated?”
- Define *E*_*a*_ as “country *A* cites on year *y*_*i*_”, *E*_*b*_ as “country *B* is cited only for its publications published on year *y*_*j*_”, and *U* as the set of total citations of publications that are published on year *y*_*j*_ and cited on *y*_*i*_. Of course, we have *y*_*i*_ ≥ *y*_*j*_. Then we can study the short-term influence if *y*_*i*_ and *y*_*j*_ are close and the long-term influence if *y*_*i*_ and *y*_*j*_ are far apart.

The 2 × 2 contingency table in Table 1 contains numbers of citations in different categories. This means the basic element is a citation in the analysis. The contingency table can also be constructed from the number of publications. In this way, *E*_*a*_ is “a publication is produced by authors from country *A*”, and *E*_*b*_ is “a publication is produced by authors from country *B*”. Then *k* in Table 1 measures the number of publications produced by authors from both *A* and *B*, i.e., via collaborations from the two countries. This implies, that the citation enrichment analysis can be extended to the publication level to study the levels of international collaborations, and eventually to construct a collaboration world map. However, it is worth noting, in the random scenario of the citation analysis, a citation can be randomly picked from the globe independently and with equal probability, as it is believed “sicence has no borders” and all publications are in theory accessible and citable. But it is different for studying international collaborations as “collaboration has borders” especially due to political policies. This may result in the world collaboration patterns being much more isolated than citation patterns.

Last, similar to the domestic citation analysis presented in this study, we can perform a journal-domestic citation analysis to study self-citations in journals. As publications in the same journal are from similar research fields^13^, it is expected there will be a high journal-domestic preference if taking total citations from the globe as the universe. Here we can restrict the universe to the citations only from the research fields of the journal to correct for such “scientific preference” of self-citations.

## CRediT authorship contribution statement

Zuguang Gu: Conceptualization; Formal analysis; Investigation; Methodology; Software; Visualization; Writing - original draft; Writing - review & editing.

## Data availability

The processed dataset is available at https://zenodo.org/doi/10.5281/zenodo.10981961. The scripts for reproducing the complete analysis in this study are available at https://github.com/jokergoo/citation_analysis.

## Acknowledgements

This research did not receive any specific grant from funding agencies in the public, commercial, or not-for-profit sectors.

## Footnotes

1 A publication can exist in both sets but marked as the citing and cited publication.

2 https://ftp.ncbi.nlm.nih.gov/pubmed/baseline/ and https://ftp.ncbi.nlm.nih.gov/pubmed/updatefiles/.

3 https://worldpopulationreview.com/country-rankings/official-names-of-countries.

4 https://github.com/w8r/k-d-tree/blob/master/data/data.tsv.

5 https://github.com/rinvex/universities/.

6 https://mjl.clarivate.com/collection-list-downloads.

7 In the remaining part of this paper, “countries and regions” are described only as countries for simplicity.

8 The average of the ranking from all cited countries and the ranking from all citing countries.

9 Domestic citations were also excluded from this matrix.

10 The value of *s* in the parentheses is the average enrichment score from the included countries. It is the same in the remaining part of the article.

11 See the explaination on the negative *R*^2^ in the caption of Table 3.

12 *−* 0.55 and *−* 0.68 are the average of the *β* and *η* coefficients of Poland, Brazil, Saudi Arabia, Thailand and mainland China on World-1.

13 For comprehensive journals or mega journals which cover several research fields, we can use “research categories” from these journals for the analysis instead.

