## Supplementary File 1 for "Two Separated Worlds: on the Preference of Influence in Life Science and Biomedical Research"

Approximately domestic publications


### Approximately domestic publications

###### Zuguang Gu

#### 2024-12-04

In current bibliometrics studies, a domestic publication is required to have all authors affiliated to the same country. However, according to my experience as a long-term biology researcher, in biology publications there are also cases where a small fraction (< 20%) of international authors exist in the middle of the author list, but this won’t affect the fact that the research conducted in the publication is domestic to that country.

In the following example (pmid: 10708385 from PubMed), there are nine authors where eight are from Spain. There is only one “international” author which is on location seven of the author list. We would assume this author contributes less than other authors and we would take this publication as a domestic publication of Spain.

If we call domestic publications where all authors are from the same country “*the strictly domestic publications*”, and publications where we allow a small fraction of international authors in the middle of the author list “*the approximately domestic publications*”. The followng plot illustrates the fraction of approximately domestic publications in different countries. On average, approximately domestic publications only contribute 5.2% to all domenstic publications.

Including the approximately domestic publications or not won’t affect the citation enrichment analysis conducted in this work. The following plot shows the log2 citation enrichments from the two settings are almost perfectly linear.
