## Supplementary File 2 for "Two Separated Worlds: on the Preference of Influence in Life Science and Biomedical Research"

Distribution of the enrichment scores


### Distribution of the enrichment scores

###### Zuguang Gu

#### 2024-12-04

The number of citations from country *A* to country *B* follows a hypergeometric distribution \(X \sim \mathrm{Hyper}(m, n, N)\) where \(m\) is the total number of times that *B* is cited, \(n\) is the total number of times that *A* cites, \(N\) is the total number of citations in the globe. To simplify the problem, we use the binomial approximation of the hypergeometric distribution: \(X \sim \mathrm{Bi}(p, n)\) where \(p = m/N\). The fold enrichment is calculated as

\[
r = \frac{k}{k\_\mathrm{exp}} = \frac{k}{np}
\]

where \(k\) is the observed number of citations and \(k\_\mathrm{exp}\) is the expected number.

Assume the observed \(k\) has a offset to the expected \(k\_\mathrm{exp}\) of \(a \cdot \sigma\) where \(\sigma\) is the standard deviation of the bimonial distribution and \(a\) is the fold factor on \(\sigma\), then

\[
k = np + a \cdot \sigma = np + a \sqrt{np(1-p)}
\]

And we can have the relation between \(r\) and \(k\) (by cancelling \(n\)) as:

\[
r = \frac{\sqrt{1 + \frac{4k}{a^2(1-p)}} + 1}{\sqrt{1 + \frac{4k}{a^2(1-p)}} - 1} = 1 + \frac{2}{\sqrt{1 + \frac{4k}{a^2(1-p)}} - 1}
\]

We can write a function which calculates \(\log\_2 r\) based on the value of \(k\):

```
f = function(k, a = 1, p = 0.5) {
    r = 1 + 2/(sqrt(1 + 4*k/a^2/(1-p)) - 1)
    log2(r)
}
```

We then demonstrate the relation between \(\log\_2 r\) and \(k\) by a simple simulation. We generate a list of random \(k\) from the binomial distribution where the size \(n\) is also randomly generated where \(\log\_{10} n\) is from a uniform distribution in the internal \([3, 7]\).

```
set.seed(123)
p = 0.1
size = round(10^(runif(10000, min = 3, max = 7)))
k = sapply(size, function(x) rbinom(1, x, p))
```

We plot \(\log\_2 r\) against \(k\), also their theoretical relations (with 1/2/3 offsets of \(\sigma\) to the expected value):

```
plot(k, log2(k/(size*p)), log = "x", pch = 16, cex = 0.5, col = "#00000080", ylab = "log2(r)")

od = order(k)
lines(k[od], f(k[od], a = 1, p = p), col = 2)
lines(k[od], f(k[od], a = 2, p = p), col = 3)
lines(k[od], f(k[od], a = 3, p = p), col = 4)

legend("topright", legend = c("k_exp + 1*sd", "k_exp + 2*sd", "k_exp + 3*sd"), lty = 1, col = 2:4)
```

So here we can see the range of the log2 fold enrichment (when the enrichment is positive) decreases when the number of citations increases.
