## Supplementary File 3 for "Two Separated Worlds: on the Preference of Influence in Life Science and Biomedical Research"

The linear fit of domestic citations


### The linear fit of domestic citations

###### Zuguang Gu

#### 2024-12-04

We first load the results of domestic citation analysis.

```
df2 = readRDS("domestic_enrichment.rds")
head(df2)
```

```
##   country_cited country_citing citations total_cited total_citing
## 1     Argentina      Argentina     33044      251784       296410
## 2     Australia      Australia    510342     3366179      2826952
## 3       Austria        Austria     86571      784203       777519
## 4    Bangladesh     Bangladesh      5626       30987        65751
## 5       Belgium        Belgium    101080     1211125       890806
## 6        Brazil         Brazil    315328     1222739      1957467
##   global_citations    p_cited   p_citing p_global_cited p_global_citing
## 1        139760456 0.11148072 0.13123948   0.0018015396     0.002120843
## 2        139760456 0.18052730 0.15160869   0.0240853464     0.020227123
## 3        139760456 0.11134262 0.11039361   0.0056110507     0.005563226
## 4        139760456 0.08556524 0.18156001   0.0002217151     0.000470455
## 5        139760456 0.11347027 0.08345959   0.0086657202     0.006373806
## 6        139760456 0.16108982 0.25788660   0.0087488195     0.014005872
##      expected  var_hyper p_hyper  log2_fc  z_score
## 1   533.99436   531.9019       0 5.951420 1409.618
## 2 68088.11826 65104.1370       0 2.905989 1733.275
## 3  4362.69850  4314.0847       0 4.310591 1251.617
## 4    14.57799    14.5679       0 8.592174 1470.193
## 5  7719.47551  7603.8047       0 3.710851 1070.651
## 6 17125.52542 16737.9380       0 4.202633 2304.943
```

We will use the following columns:

- `citations`: The number of citations that a country A cites itself, denoted as \(k\_{A-A}\).
- `total_cited`: The number of total citations that A is cited globally, denoted as \(m\_{A-}\).
- `total_citing`: The number of total citations that A cites globally, denoted as \(n\_{-A}\).
- `global_citation`: The total number of citations in the globe, denoted as \(N\).
- `expected`: The expected number of citations that A cites itself, calculated as \(m\_{A-}n\_{-A}/N\).
- `log2_fc`: The enrichment score.

Empirically, \(k\_{A-A}\) has linear relations to \(n\_{-A}\) and \(m\_{A-}\), and it can be demonstrated by the following two scatter plots (each data point corresponds to one country):

\[
\begin{align\*}
\log\_2 n\_{-A} &= a\_1 \cdot \log\_2 k\_{A-A} + b\_1 \\
\log\_2 m\_{A-} &= a\_2 \cdot \log\_2 k\_{A-A} + b\_2 \\
\end{align\*}
\]

```
par(mfrow = c(1, 2))
plot(df2$citations, df2$total_citing, log = "xy", xlab = "# Domestic citations", ylab = "# Citations A cites")
plot(df2$citations, df2$total_cited,  log = "xy", xlab = "# Domestic citations", ylab = "# Citations A is cited")
```

We fit linear regression models on the log2-scale:

```
summary(lm(log2(df2$total_citing) ~ log2(df2$citations)))
```

```
## 
## Call:
## lm(formula = log2(df2$total_citing) ~ log2(df2$citations))
## 
## Residuals:
##      Min       1Q   Median       3Q      Max 
## -0.97724 -0.26525  0.08968  0.27716  1.02866 
## 
## Coefficients:
##                     Estimate Std. Error t value Pr(>|t|)    
## (Intercept)          6.19862    0.20689   29.96   <2e-16 ***
## log2(df2$citations)  0.80860    0.01387   58.30   <2e-16 ***
## ---
## Signif. codes:  0 '***' 0.001 '**' 0.01 '*' 0.05 '.' 0.1 ' ' 1
## 
## Residual standard error: 0.4178 on 80 degrees of freedom
## Multiple R-squared:  0.977,  Adjusted R-squared:  0.9767 
## F-statistic:  3399 on 1 and 80 DF,  p-value: < 2.2e-16
```

```
summary(lm(log2(df2$total_cited) ~ log2(df2$citations)))
```

```
## 
## Call:
## lm(formula = log2(df2$total_cited) ~ log2(df2$citations))
## 
## Residuals:
##      Min       1Q   Median       3Q      Max 
## -1.91796 -0.32821 -0.05223  0.40343  1.06195 
## 
## Coefficients:
##                     Estimate Std. Error t value Pr(>|t|)    
## (Intercept)          4.44739    0.27505   16.17   <2e-16 ***
## log2(df2$citations)  0.89189    0.01844   48.37   <2e-16 ***
## ---
## Signif. codes:  0 '***' 0.001 '**' 0.01 '*' 0.05 '.' 0.1 ' ' 1
## 
## Residual standard error: 0.5554 on 80 degrees of freedom
## Multiple R-squared:  0.9669, Adjusted R-squared:  0.9665 
## F-statistic:  2340 on 1 and 80 DF,  p-value: < 2.2e-16
```

In the manuscript, we demonstrated there is a negative linear relation between the enrichment scores and the number of domestic citations:

\[
\begin{align\*}
s\_{A-A} &= \log\_2 k\_{A-A} - \log\_2 n\_{-A} + \log\_2 N - \log\_2 m\_{A-} \\
&= (1 - a\_1 - a\_2) \log\_2 k + \log\_2 N - b\_1 - b\_2 \\
&= -0.70 \log\_2 k\_{A-A} + 16.41
\end{align\*}
\]

```
plot(df2$citations, df2$log2_fc, log = "x", xlab = "# Domestic citations", ylab = "Enrichment scores")
```

```
summary(lm(df2$log2_fc ~ log2(df2$citations)))
```

```
## 
## Call:
## lm(formula = df2$log2_fc ~ log2(df2$citations))
## 
## Residuals:
##      Min       1Q   Median       3Q      Max 
## -2.09061 -0.37590 -0.08003  0.20635  2.89521 
## 
## Coefficients:
##                     Estimate Std. Error t value Pr(>|t|)    
## (Intercept)         16.41237    0.37658   43.58   <2e-16 ***
## log2(df2$citations) -0.70049    0.02525  -27.75   <2e-16 ***
## ---
## Signif. codes:  0 '***' 0.001 '**' 0.01 '*' 0.05 '.' 0.1 ' ' 1
## 
## Residual standard error: 0.7605 on 80 degrees of freedom
## Multiple R-squared:  0.9059, Adjusted R-squared:  0.9047 
## F-statistic: 769.9 on 1 and 80 DF,  p-value: < 2.2e-16
```

---

In this document, we will prove when there is a positive domestic enrichment, \(a\_1 + a\_2 > 1\), it results in the decrease of the enrichment scores.

First let’s simplify the notations: \(k\_{A-A}\) to \(k\), \(m\_{A-}\) to \(m\) and \(n\_{-A}\) to \(n\). From the two linear fits, we have:

\[
\begin{align\*}
n &= k^{a\_1} \cdot 2^{b\_1} \\
m &= k^{a\_2} \cdot 2^{b\_2} \\
mn &= k^{a\_1 + a\_2} \cdot 2^{b\_1 + b\_2} \\
\end{align\*}
\]

Denote \(k\_\mathrm{exp}\) as the expected number of citations, and \(mn\) can also be calculated as:

\[
mn = k\_\mathrm{exp}N
\]

It is easy to see, when there is no enrichment of domestic citations (i.e. \(k = k\_\mathrm{exp}\)), \(a\_1 + a\_2 = 1\).

Now we have (by cancelling \(mn\)):

\[
k^{a\_1 + a\_2} \cdot 2^{b\_1 + b\_2} = k\_\mathrm{exp}N
\]

Since all the countries share the same coefficients from the linear fit, we can simply treat different countries correspond to different stages of the citation network expansion, where small countries correspond to early states. Then for two countries with \(k\_2 > k\_1\) and \(k\_{\mathrm{exp},2} > k\_{\mathrm{exp},1}\), we divide the previous equation on both sides:

\[
\begin{align\*}
\left( \frac{k\_2}{k\_1} \right)^{a\_1 + a\_2} &= \frac{k\_{\mathrm{exp},2}}{k\_{\mathrm{exp},1}} \\
(a\_1 + a\_2)\cdot \log(k\_2/k\_1) &= \log(k\_{\mathrm{exp},2}/k\_{\mathrm{exp},1}) \\
a\_1 + a\_2 &= \frac{\log(k\_{\mathrm{exp},2}/k\_{\mathrm{exp},1})}{\log(k\_2/k\_1)} \\
\end{align\*}
\]

Denote \(T\_{A-A}\), \(T\_{A-}\) and \(T\_{-A}\) as the citation networks constructed by the citation relations in the three groups (A-domestic, A-citing and A-cited). Let’s assume from country 1 to 2, the two background networks \(T\_{A-}\) and \(T\_{-A}\) both expand by a factor of \(\alpha\) (\(\alpha > 1\)), i.e., \(m\_2 = \alpha \cdot m\_1\) and \(n\_2 = \alpha \cdot n\_1\). Then the expansion rate of the citation network if there is no enrichment is

\[
\frac{k\_{\mathrm{exp},2}}{k\_{\mathrm{exp},1}} = \frac{m\_2n\_2/N}{m\_1n\_2/N} = \alpha^2
\]

which is quadratic to \(\alpha\).

Next we look at the expansion rate of \(k\_2/k\_1\). If there is an over-representation, \(k\) can be decomposed to

\[k = k\_\mathrm{exp} + k\_\mathrm{diff}\]

where \(k\_\mathrm{diff}\) is the over-represented portion of \(k\). It can be expected that when \(m\) or \(n\) increases, \(k\_\mathrm{diff}\) also increases. Now the question is does it increase by a fixed a rate or does it still depend on \(m\) or \(n\) (i.e. increase faster or slower when \(m\) or \(n\) gets larger?)

We next check how \(k\_\mathrm{diff}/m\) or \(k\_\mathrm{diff}/n\) changes to the size of \(T\_{A-}\) and \(T\_{-A}\), i.e.

```
diff = df2$citations - df2$expected
par(mfrow = c(1, 2))
plot(df2$total_cited, diff/df2$total_cited, log = "x", xlab = "# Citations A is cited (m)", ylab = "diff/m")
plot(df2$total_citing, diff/df2$total_citing, log = "x", xlab = "# Citations A cites (n)", ylab = "diff/n")
```

We can see, in both plots, there are positive linear relations in the log-scale. Let’s fit the two linear models. Note China (index = 11), Ethiopia (index = 20) and United States (index = 80) are removed from the fitting because they are outliers on the scatter plots.

```
p1 = diff/df2$total_cited
summary(lm(p1[-c(11, 20, 80)] ~ log(df2$total_cited[-c(11, 20, 80)])))
```

```
## 
## Call:
## lm(formula = p1[-c(11, 20, 80)] ~ log(df2$total_cited[-c(11, 
##     20, 80)]))
## 
## Residuals:
##       Min        1Q    Median        3Q       Max 
## -0.080647 -0.031997 -0.009989  0.028423  0.118527 
## 
## Coefficients:
##                                      Estimate Std. Error t value Pr(>|t|)   
## (Intercept)                          0.093221   0.032033   2.910  0.00472 **
## log(df2$total_cited[-c(11, 20, 80)]) 0.003264   0.002642   1.235  0.22048   
## ---
## Signif. codes:  0 '***' 0.001 '**' 0.01 '*' 0.05 '.' 0.1 ' ' 1
## 
## Residual standard error: 0.04618 on 77 degrees of freedom
## Multiple R-squared:  0.01943,    Adjusted R-squared:  0.006697 
## F-statistic: 1.526 on 1 and 77 DF,  p-value: 0.2205
```

```
p2 = diff/df2$total_citing
summary(lm(p2[-c(11, 20, 80)] ~ log(df2$total_citing[-c(11, 20, 80)])))
```

```
## 
## Call:
## lm(formula = p2[-c(11, 20, 80)] ~ log(df2$total_citing[-c(11, 
##     20, 80)]))
## 
## Residuals:
##       Min        1Q    Median        3Q       Max 
## -0.049806 -0.018680 -0.006224  0.015745  0.078387 
## 
## Coefficients:
##                                       Estimate Std. Error t value Pr(>|t|)    
## (Intercept)                           -0.09089    0.02217  -4.099 0.000102 ***
## log(df2$total_citing[-c(11, 20, 80)])  0.01495    0.00178   8.401 1.71e-12 ***
## ---
## Signif. codes:  0 '***' 0.001 '**' 0.01 '*' 0.05 '.' 0.1 ' ' 1
## 
## Residual standard error: 0.02772 on 77 degrees of freedom
## Multiple R-squared:  0.4783, Adjusted R-squared:  0.4715 
## F-statistic: 70.58 on 1 and 77 DF,  p-value: 1.713e-12
```

To reduce the complexity of the following analysis, we assume \(m = n\).

**Scenario 1**: The first regression is not significant, then we can assume \(p = k\_\mathrm{diff}/m\) is a constant and not dependent to \(m\), i.e.:

\[
\begin{align\*}
k\_\mathrm{diff} &= pm \\
k &= k\_\mathrm{exp} + pm \\
\end{align\*}
\]

Then we can calculate \(k\_2/k\_1\) as:

\[
\begin{align\*}
\frac{k\_2}{k\_1} &= \frac{k\_{\mathrm{exp},2} + p m\_2}{k\_{\mathrm{exp},1} + p m\_1} \\
&= \frac{\alpha^2 k\_{\mathrm{exp},1} + p \alpha m\_1}{k\_{\mathrm{exp},1} + p m\_1} \\
&= \frac{\alpha^2 (k\_{\mathrm{exp},1} + p m\_1 ) + p \alpha m\_1 - p \alpha^2 m\_1}{k\_{\mathrm{exp},1} + p m\_1} \\
&= \alpha^2 - \frac{p \alpha(\alpha - 1) m\_1}{k\_{\mathrm{exp},1} + p m\_1} \\
\end{align\*}
\]

Note since \(\alpha > 1\), we have \(k\_2/k\_1 < \alpha^2\).

In the second line of above equation, it also shows the expansion of the citation network can be decomposed into two parts, the part without citation enrichment (\(\alpha^2 k\_\mathrm{exp, 1}\)) which grows in quadratic speed (\(\sim \alpha^2\)), and the “over-represented” part (\(p\alpha m\_1\)) which grows in linear speed (\(\sim \alpha\), the second line in the equation above). Mixing these two speeds results the final speed being weaker than quadratic.

**Scenario 2**: The first regression is significant. Let’s assume there is the following relation where \(p\) is dependent on \(\log m\). Note for simplicity we removed the intercept (the value of the intercept is very small and it can be ignored).

\[
p = w \log(m)
\]

where \(w\) is slop coefficient from the linear fit.

Then \(k\_2/k\_1\) can be written as:

\[
\begin{align\*}
\frac{k\_2}{k\_1} &= \frac{k\_{\mathrm{exp},2} + p\_2 \cdot m\_2}{k\_{\mathrm{exp},1} + p\_1 \cdot m\_1} \\
&= \frac{\alpha^2 \cdot k\_{\mathrm{exp},1} + (w\log (\alpha m\_1)) \cdot \alpha m\_1}{k\_{\mathrm{exp},1} + (w\log m\_1) \cdot m\_1} \\
&= \frac{\alpha^2 (k\_{\mathrm{exp},1} + w m\_1 \log m\_1 ) - \alpha^2 w m\_1\log m\_1 + \alpha w m\_1 \log \alpha + \alpha w m\_1 \log m\_1}{k\_{\mathrm{exp},1} + w m\_1 \log m\_1} \\
&= \alpha^2 - \frac{\alpha w m\_1 \left[ (\alpha - 1) \log m\_1 - \log \alpha \right]}{k\_{\mathrm{exp},1} + w m\_1 \log m\_1} \\
\end{align\*}
\]

where \(\log m\_1\) is much larger than \(\log \alpha\), which also results in \(k\_2/k\_1 < \alpha^2\).

In this case, in the expansion of the citation network, the “over-represented” part grows in a speed of \(\sim \alpha \log \alpha\) (the second line in above equation), but still slower than quadratic.

For both scenarios, we can have:

\[
\begin{align\*}
k\_{\mathrm{exp},2}/ k\_{\mathrm{exp},1} &> k\_2 / k\_1 > 1 \\
\log(k\_{\mathrm{exp},2}/ k\_{\mathrm{exp},1}) &> \log( k\_2 / k\_1 ) > 0 \\
\end{align\*}
\]

And eventually

\[
a\_1 + a\_2 = \frac{\log(k\_{\mathrm{exp},2}/k\_{\mathrm{exp},1})}{\log(k\_2/k\_1)} > 1
\]
