## Supplementary File 4 for "Two Separated Worlds: on the Preference of Influence in Life Science and Biomedical Research"

Visualize the grouping patterns of countries


### Visualize the grouping patterns of countries

###### Zuguang Gu

#### 2024-12-04

- Dimension reduction by MDS, t-SNE and UMAP
- Robustness of the partitioning

#### Dimension reduction by MDS, t-SNE and UMAP

In the manuscript, countries are clustered into 7 groups based on their preference of influence patterns by a graph clustering method. In this document, we will apply dimension reduction methods to validate the clustering results.

The influence matrix contains `NA` values where two countries do not have enough number of citations for calculating the influence value. Also there might be “outliers” when calculating the distance between two countries. Thus, we first defined a “robust distance function” which removes `NA` values and also extreme values.

```
robust_dist_pair = function(x, y, trim = 0.1) {
    x_mid = mean(x, na.rm = TRUE)
    y_mid = mean(y, na.rm = TRUE)

    dd = sqrt( (x-x_mid)^2 + (y-y_mid)^2 )

    q = quantile(dd, c(trim/2, 1-trim/2), na.rm = TRUE)
    l = dd >= q[1] & dd <= q[2]
    l[is.na(l)] = FALSE
    sqrt( sum( (x[l] - y[l])^2 ) )/sum(l)
}

# by rows
robust_dist = function(m, trim = 0.1) {
    n = nrow(m)
    d = matrix(0, nrow = n, ncol = n)
    rownames(d) = colnames(d) = rownames(m)

    for(i in 1:(n-1)) {
        for(j in (i+1):n) {
            d[i, j] = d[j, i] = robust_dist_pair(m[i, ], m[j, ], trim = trim)
        }
    }

    as.dist(d)
}
```

We load the data. `m` is the influence matrix of 72 countries. `mem` is the clustering results from the graph clustering analysis.

```
m = readRDS("mat_influence.rds")
mem = readRDS("country_groups.rds")

identical(rownames(m), names(mem))
```

```
## [1] TRUE
```

```
identical(colnames(m), names(mem))
```

```
## [1] TRUE
```

```
split(names(mem), mem)
```

```
## $`1`
## [1] "Argentina" "Brazil"    "Chile"     "Colombia"  "Mexico"    "Peru"     
## [7] "Uruguay"  
## 
## $`2`
##  [1] "Australia"      "Austria"        "Belgium"        "Canada"        
##  [5] "Denmark"        "Estonia"        "Finland"        "France"        
##  [9] "Germany"        "Ireland"        "Israel"         "Luxembourg"    
## [13] "Netherlands"    "New Zealand"    "Norway"         "Sweden"        
## [17] "Switzerland"    "United Kingdom"
## 
## $`3`
##  [1] "Bangladesh"  "China"       "Hong Kong"   "India"       "Indonesia"  
##  [6] "Japan"       "Malaysia"    "Pakistan"    "Singapore"   "South Korea"
## [11] "Taiwan"      "Thailand"    "Vietnam"    
## 
## $`4`
##  [1] "Bulgaria"       "Croatia"        "Czech Republic" "Greece"        
##  [5] "Hungary"        "Italy"          "Lithuania"      "Poland"        
##  [9] "Portugal"       "Romania"        "Russia"         "Serbia"        
## [13] "Slovakia"       "Slovenia"       "Spain"         
## 
## $`5`
##  [1] "Egypt"                "Iran"                 "Jordan"              
##  [4] "Kuwait"               "Lebanon"              "Morocco"             
##  [7] "Qatar"                "Saudi Arabia"         "Tunisia"             
## [10] "Turkey"               "United Arab Emirates"
## 
## $`6`
## [1] "Ethiopia"     "Ghana"        "Kenya"        "Nigeria"      "South Africa"
## [6] "Tanzania"     "Uganda"      
## 
## $`7`
## [1] "United States"
```

We will try the following dimension reduction methods: MDS (multidimensional scaling), UMAP and t-SNE. For each method, the distance will be calculated on rows and columns of `m` separately, thus there are two plots for each method.

```
library(ggplot2)
library(ggrepel)
library(cowplot)
library(grid)
```

```
d = robust_dist(m)
loc = cmdscale(d)

loc = as.data.frame(loc)[, 1:2]
colnames(loc) = c("V1", "V2")
loc$label = names(mem)
loc$group = paste0("group-", mem)

p1 = ggplot(loc, aes(x = V1, y = V2, color = group, label = label)) + 
    geom_point() + geom_text_repel(max.overlaps = Inf, size = 3, show.legend = FALSE) +
    labs(x = "MDS 1", y = "MDS 2") +
    ggtitle("As cited countries / how they influence other countries")
p1 = grid.grabExpr(print(p1))


# the second plot, note `m` is transposed
d = robust_dist(t(m))
loc = cmdscale(d)

loc = as.data.frame(loc)[, 1:2]
colnames(loc) = c("V1", "V2")
loc$label = names(mem)
loc$group = paste0("group-", mem)

p2 = ggplot(loc, aes(x = V1, y = V2, color = group, label = label)) + 
    geom_point() + geom_text_repel(max.overlaps = Inf, size = 3, show.legend = FALSE) +
    labs(x = "MDS 1", y = "MDS 2") +
    ggtitle("As citing countries / how they are influenced by other countries")
p2 = grid.grabExpr(print(p2))

plot_grid(p1, p2, nrow = 1)
```

Next we apply UMAP and t-SNE dimension reduction methods. Note they need a complete matrix with no `NA` value. Thus, we first impute the `NA` values with the **impute** package.

```
library(impute)
m2 = impute.knn(m)$data
```

We apply a wrapper function `dimension_reduction()` from the **cola** package for UMAP and t-SNE analysis. Note `dimension_reduction()` is applied on columns.

```
library(cola)
set.seed(123)
# on the rows of `m2`
loc = dimension_reduction(t(m2), method = "UMAP")

loc = as.data.frame(loc)[, 1:2]
colnames(loc) = c("V1", "V2")
loc$label = names(mem)
loc$group = paste0("group-", mem)
```

```
p1 = ggplot(loc, aes(x = V1, y = V2, color = group, label = label)) + 
    geom_point() + geom_text_repel(max.overlaps = Inf, size = 3, show.legend = FALSE) +
    labs(x = "UMAP 1", y = "UMAP 2") +
    ggtitle("As cited countries / how they influence other countries")
p1 = grid.grabExpr(print(p1))
```

```
# on the columns of m2
loc = dimension_reduction(m2, method = "UMAP")
loc = as.data.frame(loc)[, 1:2]
colnames(loc) = c("V1", "V2")
loc$label = names(mem)
loc$group = paste0("group-", mem)
```

```
p2 = ggplot(loc, aes(x = V1, y = V2, color = group, label = label)) + 
    geom_point() + geom_text_repel(max.overlaps = Inf, size = 3, show.legend = FALSE) +
    labs(x = "UMAP 1", y = "UMAP 2") +
    ggtitle("As citing countries / how they are influenced by other countries")
p2 = grid.grabExpr(print(p2))

plot_grid(p1, p2, nrow = 1)
```

And the dimension reduction by t-SNE.

```
# on the rows of `m2`
loc = dimension_reduction(t(m2), method = "t-SNE")

loc = as.data.frame(loc)[, 1:2]
colnames(loc) = c("V1", "V2")
loc$label = names(mem)
loc$group = paste0("group-", mem)
```

```
p1 = ggplot(loc, aes(x = V1, y = V2, color = group, label = label)) + 
    geom_point() + geom_text_repel(max.overlaps = Inf, size = 3, show.legend = FALSE) +
    labs(x = "t-SNE 1", y = "t-SNE 2") +
    ggtitle("As cited countries / how they influence other countries")
```

```
# on the columns of `m2`
loc = dimension_reduction(m2, method = "t-SNE")

loc = as.data.frame(loc)[, 1:2]
colnames(loc) = c("V1", "V2")
loc$label = names(mem)
loc$group = paste0("group-", mem)
```

```
p2 = ggplot(loc, aes(x = V1, y = V2, color = group, label = label)) + 
    geom_point() + geom_text_repel(max.overlaps = Inf, size = 3, show.legend = FALSE) +
    labs(x = "t-SNE 1", y = "t-SNE 2") +
    ggtitle("As citing countries / how they are influenced by other countries")
p2 = grid.grabExpr(print(p2))

plot_grid(p1, p2, nrow = 1)
```

#### Robustness of the partitioning

There are two steps when performing the graph-clustering method:

1. constructing an undirected graph. When the values for both `A->B` and `B->A` are both available, there are three modes for aggregating the two values into one single value for `A-B`. In the graph construction function `graph_from_adjacency_matrix()`, we can set `"max"`, `"min"` or `"plus"` to the `mode` argument which corresponds to taking the minimal, the maximal or the sum of the two directional values.
2. in clusteing, there is a parameter `resolution` which controls the “tightness” of the links within clusters.

We try all three values of `mode` and a list of cutoffs for `resolution`, and we evaluate the robustness of the partitioning.

Note the classification of 1-7 that splits the heatmap columns (the country groups) is from the manuscript.

```
m2[m2 < 0.0001] = 0
m2[is.na(m2)] = 0
library(igraph)

ml = list()
for(mode in c("max", "min", "plus")) {

    g = graph_from_adjacency_matrix(m2, mode = mode, weighted = TRUE)
    for(resolution in seq(1.1, 1.3, by = 0.05)) {
        cm = cluster_louvain(g, weight = E(g)$weight, resolution = resolution)
        communities(cm)
        ml[[paste0(mode, "-", resolution)]] = membership(cm)
    }
}

ml = lapply(ml, function(x) cola:::relabel_class(x, mem, return_map = FALSE))
mm = do.call(rbind, ml)
colnames(mm) = names(mem)

library(ComplexHeatmap)
Heatmap(mm, name = "Classifications", column_split = mem, 
    col = structure(RColorBrewer::brewer.pal(9, "Set1"), names = 1:9), 
    cluster_column_slices = FALSE,
    show_row_dend = FALSE, show_column_dend = FALSE)
```
