## Supplementary File 5 for "Two Separated Worlds: on the Preference of Influence in Life Science and Biomedical Research"

Mutual influence between West-1 and World-2


### Mutual influence between West-1 and World-2

###### Zuguang Gu

#### 2024-05-07

We load the data. `m` is the influence matrix of 72 countries. `mem` is the clustering results from the graph clustering analysis.

```
library(ComplexHeatmap)
library(circlize)

m = readRDS("mat_influence.rds")
mem = readRDS("country_groups.rds")

lt = split(names(mem), mem)
lt
```

```
## $`1`
## [1] "Argentina" "Brazil"    "Chile"     "Colombia"  "Mexico"    "Peru"     
## [7] "Uruguay"  
## 
## $`2`
##  [1] "Australia"      "Austria"        "Belgium"        "Canada"        
##  [5] "Denmark"        "Estonia"        "Finland"        "France"        
##  [9] "Germany"        "Ireland"        "Israel"         "Luxembourg"    
## [13] "Netherlands"    "New Zealand"    "Norway"         "Sweden"        
## [17] "Switzerland"    "United Kingdom"
## 
## $`3`
##  [1] "Bangladesh"  "China"       "Hong Kong"   "India"       "Indonesia"  
##  [6] "Japan"       "Malaysia"    "Pakistan"    "Singapore"   "South Korea"
## [11] "Taiwan"      "Thailand"    "Vietnam"    
## 
## $`4`
##  [1] "Bulgaria"       "Croatia"        "Czech Republic" "Greece"        
##  [5] "Hungary"        "Italy"          "Lithuania"      "Poland"        
##  [9] "Portugal"       "Romania"        "Russia"         "Serbia"        
## [13] "Slovakia"       "Slovenia"       "Spain"         
## 
## $`5`
##  [1] "Egypt"                "Iran"                 "Jordan"              
##  [4] "Kuwait"               "Lebanon"              "Morocco"             
##  [7] "Qatar"                "Saudi Arabia"         "Tunisia"             
## [10] "Turkey"               "United Arab Emirates"
## 
## $`6`
## [1] "Ethiopia"     "Ghana"        "Kenya"        "Nigeria"      "South Africa"
## [6] "Tanzania"     "Uganda"      
## 
## $`7`
## [1] "United States"
```

To be the same as in the manuscript, we extract countries in West-1 (group-1), and perform the Louvain graph clustering method to partition West-1 into three subgroups:

```
cn = lt[["2"]]
subm = m[cn, cn]
subm2 = subm
subm2[is.na(subm2)] = 0
subm2[subm2 < 0] = 0
library(igraph)
g2 = graph_from_adjacency_matrix(subm2, mode = "plus", weighted = TRUE)

set.seed(666)
mem2 = membership(cluster_louvain(g2, weight = sqrt(E(g2)$weight), resolution = 1.1))
mem2 = factor(mem2, levels = c(3, 1, 2))
levels(mem2) = c("Northern Europe", "English-speaking", "Mainland Europe")
mem2
```

```
##        Australia          Austria          Belgium           Canada 
## English-speaking  Mainland Europe  Mainland Europe English-speaking 
##          Denmark          Estonia          Finland           France 
##  Northern Europe  Northern Europe  Northern Europe  Mainland Europe 
##          Germany          Ireland           Israel       Luxembourg 
##  Mainland Europe English-speaking  Mainland Europe  Mainland Europe 
##      Netherlands      New Zealand           Norway           Sweden 
##  Mainland Europe English-speaking  Northern Europe  Northern Europe 
##      Switzerland   United Kingdom 
##  Mainland Europe English-speaking 
## Levels: Northern Europe English-speaking Mainland Europe
```

And the heatmap the same as Figure 7 in the main manuscript:

```
col_fun = colorRamp2(c(-2, 0, 2), c("blue", "white", "red"))
ht = Heatmap(subm, name = "Enrichment score",
    row_title = "Cited/influencing countries",
    column_title = "Citing/influenced countries", column_title_side = "bottom",
    col = col_fun,
    row_split = mem2, column_split = mem2,
    na_col = "#DDDDDD",
    cluster_row_slices = FALSE, cluster_column_slices = FALSE,
    show_row_dend = FALSE, show_column_dend = FALSE,
    border = "black",
    width = unit(4, "mm")*nrow(subm),
    height = unit(4, "mm")*ncol(subm)
)

draw(ht, column_title = "Influence map only in West-1 countries",
    heatmap_legend_list = list(Legend(at = "Not available", legend_gp = gpar(fill = "#DDDDDD"))))
```

We remove Estonia and Luxembourg because most of the enrichment values are missing between they two and other countries in World-2.

```
cn = setdiff(cn, c("Estonia", "Luxembourg"))
mem2 = mem2[!names(mem2) %in% c("Estonia", "Luxembourg")]
mem2_col = c("Northern Europe" = 2, "English-speaking" = 3, "Mainland Europe" = 4)
```

The first heatmap shows how World-1 countries influence World-2.

```
cn2 = setdiff(names(mem), c(lt[["2"]], "United States"))
Heatmap(m[cn, cn2], name = "Enrichment score",
    col = col_fun,
    right_annotation = rowAnnotation(group = mem2, col = list(group = mem2_col)),
    column_title = "How West-1 influences World-2",
    column_split = mem[cn2]
)
```

The second heatmap shows how West-1 is influenced by World-2.

```
draw(Heatmap(m[cn2, cn], name = "Enrichment score",
    col = col_fun,
    top_annotation = HeatmapAnnotation(group = mem2, col = list(group = mem2_col)),
    column_title = "How West-1 is influenced by World-2",
    row_split = mem[cn2]
), merge_legends = TRUE)
```

The classification of “Northern Europe/English-speaking/Mainland Europe” cannot be observed in both heatmaps.
