## Supplementary File 6 for "Two Separated Worlds: on the Preference of Influence in Life Science and Biomedical Research"

Other enrichment methods


### Other enrichment methods

###### Zuguang Gu

#### 2024-05-07

We calculate enrichment scores and partition countries.

```
source("lib.R")

df = read.table("num_cite_country_country.tab", quote = "", sep = "\t", header = TRUE)
df = calc_stat(df, cutoff_total = 10000, cutoff_single = 100, all = TRUE)

m = readRDS("mat_influence.rds")
mem = readRDS("country_groups.rds")


mem_name = mem
mem_name[mem == 1] = "Latin-America"
mem_name[mem == 2] = "West-1"
mem_name[mem == 3] = "Asia-2"
mem_name[mem == 4] = "Europe-3"
mem_name[mem == 5] = "Mid-East"
mem_name[mem == 6] = "Africa"
mem_name[mem == 7] = "US"

mem_name["Hong Kong"] = "Asia-1"
mem_name["Singapore"] = "Asia-1"
mem_name["Taiwan"] = "Asia-1"
mem_name["South Korea"] = "Asia-1"
mem_name["China"] = "Asia-1"
mem_name["Japan"] = "Asia-1"

mem_name["Spain"] = "Europe-2"
mem_name["Italy"] = "Europe-2"
mem_color = structure(RColorBrewer::brewer.pal(length(setdiff(unique(mem_name), NA)), "Set1"), names = setdiff(unique(mem_name), NA))
mem_color2 = mem_color[setdiff(names(mem_color), "Africa")]

plot_balance = function(country, group, ...) {
    
    df1 = df[df$country_cited %in% lt[[group]] & df$country_citing != df$country_cited & !df$country_citing %in% names(mem_name)[mem_name == "Africa"], ]
    df2 = df[df$country_citing %in% lt[[group]] & df$country_citing != df$country_cited& !df$country_cited %in% names(mem_name)[mem_name == "Africa"], ]
    rg = range(c(df1$log2_fc, df2$log2_fc))

    df1 = df[df$country_citing == country & df$country_cited != country, ]
    df2 = df[df$country_cited == country & df$country_citing != country, ]

    rownames(df1) = df1$country_cited
    rownames(df2) = df2$country_citing
    cn = intersect(rownames(df1), rownames(df2))
    cn = setdiff(cn, names(mem_name)[mem_name == "Africa"])

    df1 = df1[cn, ]
    df2 = df2[cn, ]

    
    plot(df1$log2_fc, df2$log2_fc, pch = 16, col = mem_color[mem_name[rownames(df1)]], 
        xlab = "s_{B-A}", ylab = "s_{A-B}", main = paste0(country, " (country A)"), 
        xlim = rg, ylim = rg, ...)
    abline(h = 0, col = "grey")
    abline(v = 0, col = "grey")
    abline(a = 0, b = 1, col = "grey", lty = 2)
}

lt = split(names(mem_name), mem_name)
```

Then we plot mutual influence for individual countries in different country-groups.

##### Asia-1

```
par(mfrow = c(2, 4))
for(country in lt[["Asia-1"]]) {
    plot_balance(country, "Asia-1")
}
plot.new()
legend("left", legend = names(mem_color2), pch = 16, col = mem_color2)
```

##### Asia-2

```
par(mfrow = c(2, 4))
for(country in lt[["Asia-2"]]) {
    plot_balance(country, "Asia-2")
}
plot.new()
legend("left", legend = names(mem_color2), pch = 16, col = mem_color2)
```

##### Europe-2

```
par(mfrow = c(1, 4))
for(country in lt[["Europe-2"]]) {
    plot_balance(country, "Europe-2")
}
plot.new()
legend("left", legend = names(mem_color2), pch = 16, col = mem_color2)
plot.new()
```

##### Europe-3

```
par(mfrow = c(4, 4))
for(country in lt[["Europe-3"]]) {
    plot_balance(country, "Europe-3")
}
```

##### Latin-America

```
par(mfrow = c(2, 4))
for(country in lt[["Latin-America"]]) {
    plot_balance(country, "Latin-America")
}
plot.new()
legend("left", legend = names(mem_color2), pch = 16, col = mem_color2)
```

##### Mid-East

```
par(mfrow = c(3, 4))
for(country in lt[["Mid-East"]]) {
    plot_balance(country, "Mid-East")
}
plot.new()
legend("left", legend = names(mem_color2), pch = 16, col = mem_color2)
```

##### West-1

```
par(mfrow = c(5, 4))
for(country in lt[["West-1"]]) {
    plot_balance(country, "West-1")
}
plot.new()
legend("left", legend = names(mem_color2), pch = 16, col = mem_color2)
```

### US

```
par(mfrow = c(1, 4))
for(country in lt[["US"]]) {
    plot_balance(country, "US")
}
plot.new()
legend("left", legend = names(mem_color2), pch = 16, col = mem_color2)
plot.new()
plot.new()
```
