## Supplementary File 7 for "Two Separated Worlds: on the Preference of Influence in Life Science and Biomedical Research"

Citation balance of China


### Citation balance of China

###### Zuguang Gu

#### 2024-11-22

`m` is the mutual influence matrix. `mem` is the partitioning result of countries in `m`.

```
m = readRDS("mat_influence.rds")
mem = readRDS("country_groups.rds")

mem_name = mem
mem_name[mem == 1] = "Latin-America"
mem_name[mem == 2] = "West-1"
mem_name[mem == 3] = "Asia-2"
mem_name[mem == 4] = "Europe-3"
mem_name[mem == 5] = "Mid-East"
mem_name[mem == 6] = "Africa"
mem_name[mem == 7] = "US"

mem_name["Hong Kong"] = "Asia-1"
mem_name["Singapore"] = "Asia-1"
mem_name["Taiwan"] = "Asia-1"
mem_name["South Korea"] = "Asia-1"
mem_name["China"] = "Asia-1"
mem_name["Japan"] = "Asia-1"

mem_name["Spain"] = "Europe-2"
mem_name["Italy"] = "Europe-2"
mem_color = structure(RColorBrewer::brewer.pal(length(setdiff(unique(mem_name), NA)), "Set1"), names = setdiff(unique(mem_name), NA))
```

Obtain the enrichment scores on China and from China, and remove Africa from the analysis.

```
x1 = m[, "China"]
x2 = m["China", ]

x2 = x2[names(x1)]

l = x1 != 0 & x2 != 0 & !is.na(x1) & !is.na(x2) & (names(x1) %in% names(mem_name[mem_name != "Africa"]))
x1 = x1[l]
x2 = x2[l]
```

We take \(y=x\) as the baseline for the linear fit.

```
rotate_lm = function(y, x) {
    fit = lm(y ~ x)

    d = cooks.distance(fit)
    l = d <= 0.5

    x = x[l]
    y = y[l]

    y2 = y - x
    x2 = y + x

    fit = lm(y2 ~ x2)

    slop = (1+fit$coefficients[2])/(1-fit$coefficients[2])
    intercept = fit$coefficients[1]/(1-fit$coefficients[2])

    r2 = 1 - sum((y - (intercept + slop*(x)))^2)/sum( (y - mean(y))^2 )

    list(coefficients = c(intercept, slop), r2 = r2)
}
```

Then for World-1, Asia-1, Mid-East and Europe-3, we fit a linear regression model on each country group. Asia-2 is removed because there is no reliable fit.

```
max = 2
pushViewport(viewport(xscale = c(-max, max), yscale = c(-max, max), width = unit(1, "npc") - unit(1, "cm"), height = unit(1, "npc") - unit(1, "cm")))
grid.rect(gp = gpar(fill = "#EEEEEE", col = NA))
grid.lines(c(-max, max), c(-max, max), default.units = "native", gp = gpar(col = "grey"))
grid.lines(c(0, 0), c(-max, max), default.units = "native", gp = gpar(col = "grey", lty = 2))
grid.lines(c(-max, max), c(0, 0), default.units = "native", gp = gpar(col = "grey", lty = 2))
grid.points(x1, x2, gp = gpar(col = mem_color[mem_name[names(x1)]]), pch = 16, size = unit(4, "pt"), default.units = "native")

l = mem_name[names(x1)] %in% c("West-1", "US")  # world-1
fit = rotate_lm(x2[l], x1[l])
fit
```

```
## $coefficients
## (Intercept)          x2 
##  -0.5362685   2.0354510 
## 
## $r2
## [1] 0.8602667
```

```
slop = fit$coefficients[2]
intercept = fit$coefficients[1]
grid.lines(range(x1[l]), intercept + slop*range(x1[l]), default.units = "native", gp = gpar(col = mem_color["West-1"], lwd = 2))

l = mem_name[names(x1)] %in% "Asia-1"  # Asia-1
fit = rotate_lm(x2[l], x1[l])
fit
```

```
## $coefficients
## (Intercept)          x2 
##   -1.176255    1.986670 
## 
## $r2
## [1] 0.9315182
```

```
slop = fit$coefficients[2]
intercept = fit$coefficients[1]
grid.lines(range(x1[l]), intercept + slop*range(x1[l]), default.units = "native", gp = gpar(col = mem_color["Asia-1"], lwd = 2))

l = mem_name[names(x1)] %in% "Mid-East"  # Mid-East
fit = rotate_lm(x2[l], x1[l])
fit
```

```
## $coefficients
## (Intercept)          x2 
##  -0.1228599   1.9298406 
## 
## $r2
## [1] -0.1756173
```

```
slop = fit$coefficients[2]
intercept = fit$coefficients[1]
grid.lines(range(x1[l]), intercept + slop*range(x1[l]), default.units = "native", gp = gpar(col = mem_color["Mid-East"], lwd = 2))

l = mem_name[names(x1)] %in% "Europe-3"  # Europe-3
fit = rotate_lm(x2[l], x1[l])
fit
```

```
## $coefficients
## (Intercept)          x2 
##   0.1877229   2.9555156 
## 
## $r2
## [1] 0.01384803
```

```
slop = fit$coefficients[2]
intercept = fit$coefficients[1]
grid.lines(range(x1[l]), intercept + slop*range(x1[l]), default.units = "native", gp = gpar(col = mem_color["Europe-3"], lwd = 2))

library(ComplexHeatmap)
lgd = Legend(title = "Region", at = c("World-1", "Asia-1", "Mid-East", "Europe-3"),
    type = "lines", legend_gp = gpar(col = mem_color[c(c("West-1", "Asia-1", "Mid-East", "Europe-3"))]))
draw(lgd, x = unit(-1.5, "native"), y = unit(1.5, "native"), just = c("top", "left"))

popViewport()
```

And the four linear models are:

|  |  |
| --- | --- |
| World-1 | \(y = 2.035x - 0.536\) |
| Asia-1 | \(y = 1.987x - 1.176\) |
| Mid-East | \(y = 1.930x - 0.123\) |
| Europe-3 | \(y = 2.956x + 0.188\) |
