## Supplementary File 8 for "Two Separated Worlds: on the Preference of Influence in Life Science and Biomedical Research"

Enrichment measured by z-scores


### Enrichment measured by z-scores

###### Zuguang Gu

#### 2024-05-07

```
country_meta = read.table("country_meta.tab", sep = "\t", quote = "", header = TRUE)
country_meta = country_meta[!duplicated(country_meta$standardName), ]
country_meta$subregion2 = ifelse(grepl(", ", country_meta$subregion), gsub("^.*, ", "", country_meta$subregion), NA)
country_meta$subregion = gsub(", .*$", "", country_meta$subregion)
rownames(country_meta) = country_meta$standardName
country_meta = country_meta[, c("officialName", "region", "subregion", "subregion2")]
head(country_meta)
```

```
##                               officialName        region          subregion
## India                    Republic of India          Asia      Southern Asia
## China           People's Republic of China          Asia       Eastern Asia
## United States     United States of America North America   Northern America
## Indonesia            Republic of Indonesia          Asia South-Eastern Asia
## Pakistan      Islamic Republic of Pakistan          Asia      Southern Asia
## Nigeria        Federal Republic of Nigeria        Africa     Western Africa
##                       subregion2
## India         South Central Asia
## China                       <NA>
## United States               <NA>
## Indonesia                   <NA>
## Pakistan      South Central Asia
## Nigeria       Sub-Saharan Africa
```

```
df = read.table("num_cite_country_country.tab", quote = "", sep = "\t", header = TRUE)
n_cited = tapply(df[, 3], df[, 1], sum)
n_citing = tapply(df[, 3], df[, 2], sum)

source("lib.R")

df = read.table("num_cite_country_country.tab", quote = "", sep = "\t", header = TRUE)
df = calc_stat(df, cutoff_total = 10000, cutoff_single = 100, all = TRUE)
df$cate = ifelse(df$country_cited == df$country_citing, "Domestic", 
             ifelse(country_meta[df$country_cited, "region"]=="Africa" & country_meta[df$country_citing, "region"]== "Africa", 
                "International\n  within Africa", "International"))
df$cate = factor(df$cate, levels = c("Domestic", "International\n  within Africa", "International"))
```

```
ggplot(df, aes(x = citations, y = z_score, col = cate)) + geom_point(size = 0.5) +
    scale_color_discrete() + scale_x_log10()
```

Use the same graph clustering method as in the main manuscript.

```
df = read.table("num_cite_country_country.tab", quote = "", sep = "\t", header = TRUE)
df = calc_stat(df, cutoff_total = 10000, cutoff_single = 100, min_country = 25, all = FALSE)
m = xtabs(z_score ~ country_cited + country_citing, data = df)
class(m) = "matrix"

all_countries = rownames(m)

m2 = m
m2[m2 < 0] = 0
library(igraph)

set.seed(123)
g = graph_from_adjacency_matrix(m2, mode = "plus", weighted = TRUE)
cm = cluster_louvain(g, weight = E(g)$weight, resolution = 1.2)
communities(cm)
```

```
## $`1`
## [1] "Argentina" "Brazil"    "Chile"     "Colombia"  "Mexico"    "Peru"     
## [7] "Uruguay"  
## 
## $`2`
##  [1] "Australia"      "Austria"        "Belgium"        "Canada"        
##  [5] "Denmark"        "Estonia"        "Finland"        "France"        
##  [9] "Germany"        "Ireland"        "Israel"         "Luxembourg"    
## [13] "Netherlands"    "New Zealand"    "Norway"         "Sweden"        
## [17] "Switzerland"    "United Kingdom"
## 
## $`3`
## [1] "Bangladesh"   "Ethiopia"     "Ghana"        "Kenya"        "Nigeria"     
## [6] "South Africa" "Tanzania"     "Uganda"      
## 
## $`4`
##  [1] "Bulgaria"       "Croatia"        "Czech Republic" "Greece"        
##  [5] "Hungary"        "Italy"          "Lithuania"      "Poland"        
##  [9] "Portugal"       "Romania"        "Russia"         "Serbia"        
## [13] "Slovakia"       "Slovenia"       "Spain"         
## 
## $`5`
##  [1] "China"       "Hong Kong"   "Indonesia"   "Japan"       "Malaysia"   
##  [6] "Singapore"   "South Korea" "Taiwan"      "Thailand"    "Vietnam"    
## 
## $`6`
##  [1] "Egypt"                "India"                "Iran"                
##  [4] "Jordan"               "Kuwait"               "Lebanon"             
##  [7] "Morocco"              "Pakistan"             "Qatar"               
## [10] "Saudi Arabia"         "Tunisia"              "Turkey"              
## [13] "United Arab Emirates"
## 
## $`7`
## [1] "United States"
```

```
mem = membership(cm)

region_color = structure(RColorBrewer::brewer.pal(length(unique(country_meta$region)), "Set3"), names = unique(country_meta$region))

library(ComplexHeatmap)

robust_dist_pair = function(x, y, trim = 0.1) {
    x_mid = mean(x, na.rm = TRUE)
    y_mid = mean(y, na.rm = TRUE)

    dd = sqrt( (x-x_mid)^2 + (y-y_mid)^2 )

    q = quantile(dd, c(trim/2, 1-trim/2), na.rm = TRUE)
    l = dd >= q[1] & dd <= q[2]
    l[is.na(l)] = FALSE
    sqrt( sum( (x[l] - y[l])^2 ) )/sum(l)
}

robust_dist = function(m, trim = 0.1) {
    n = nrow(m)
    d = matrix(0, nrow = n, ncol = n)
    rownames(d) = colnames(d) = rownames(m)

    for(i in 1:(n-1)) {
        for(j in (i+1):n) {
            d[i, j] = d[j, i] = robust_dist_pair(m[i, ], m[j, ], trim = trim)
        }
    }

    as.dist(d)
}

m_with_NA = m
m_with_NA[m == 0] = NA

row_order = order.dendrogram(reorder(as.dendrogram(hclust(robust_dist(m_with_NA))), -rowMeans(m_with_NA, na.rm = TRUE)))
col_order = order.dendrogram(reorder(as.dendrogram(hclust(robust_dist(t(m_with_NA)))), -colMeans(m_with_NA, na.rm = TRUE)))

ht = Heatmap(m, name = "Enrichment score", #cluster_rows = hc, cluster_columns = hc,
    row_title = "Cited/influencing countries",
    column_title = "Citing/influenced countries", column_title_side = "bottom",
    row_order = row_order, column_order = col_order, row_names_side = "left",
    row_names_gp = gpar(fontsize = 8),
    column_names_gp = gpar(fontsize = 8),
    layer_fun = function(j, i, x, y, w, h, fill) {
        l = pindex(m, i, j) == 0
        if(any(l)) grid.rect(x[l], y[l], w[l], h[l], gp = gpar(fill = "#DDDDDD", col = "#DDDDDD"))
    },
    right_annotation = rowAnnotation(Region = country_meta[rownames(m), "region"], 
                                     "#cited" = anno_barplot(sqrt(as.vector(n_cited[rownames(m)])), ylim = c(0, 8000), axis_param = list(at = c(0,2000,4000,6000,8000), labels = c("0", "4M", "16M", "36M", "64M"))),
                                     col = list(Region = region_color)),
    top_annotation = HeatmapAnnotation(Region = country_meta[colnames(m), "region"], 
                                       "#citing" = anno_barplot(sqrt(as.vector(n_citing[colnames(m)])), ylim = c(0, 8000), axis_param = list(at = c(0,2000,4000,6000,8000), labels = c("0", "4M", "16M", "36M", "64M"))),
                                       col = list(Region = region_color)),
    cluster_row_slices = FALSE, cluster_column_slices = FALSE,
    row_split = factor(as.character(mem[rownames(m)]), levels = c(2, 7, 3, 5, 6, 1, 4)),
    column_split = factor(as.character(mem[colnames(m)]), levels = c(2, 7, 3, 5, 6, 1, 4)),
    border = "black"

)
draw(ht, merge_legend = TRUE, column_title = "Enrichment of citations between countries",
    heatmap_legend_list = list(Legend(at = "Not available", legend_gp = gpar(fill = "#DDDDDD"))))
```

We compare the country classifications from log2 fold enrichment and *z*-score:

```
mem_by_log2_fc = readRDS("country_groups.rds")
mem_by_log2_fc = cola:::relabel_class(mem_by_log2_fc, mem, return_map = FALSE)
mm = rbind(as.character(mem), as.character(mem_by_log2_fc))
colnames(mm) = names(mem)
Heatmap(mm, name = "classification",
    top_annotation = HeatmapAnnotation(Region = country_meta[rownames(m), "region"], 
                                     col = list(Region = region_color)),
    column_order = order(mem, mem_by_log2_fc, country_meta[rownames(m), "region"]),
    col = structure(names = 1:7, 2:8),
    row_labels = c("by_z_score", "by_log2_fc"))
```

And the agreement is:

```
sum(mem == mem_by_log2_fc)/length(mem)
```

```
## [1] 0.9583333
```
