## Supplementary File 9 for "Two Separated Worlds: on the Preference of Influence in Life Science and Biomedical Research"

Select universe


### Select universe

###### Zuguang Gu

#### 2024-05-16

```
source("lib.R")
```

```
country_meta = read.table("country_meta.tab", sep = "\t", quote = "", header = TRUE)
country_meta = country_meta[!duplicated(country_meta$standardName), ]
country_meta$subregion2 = ifelse(grepl(", ", country_meta$subregion), gsub("^.*, ", "", country_meta$subregion), NA)
country_meta$subregion = gsub(", .*$", "", country_meta$subregion)
rownames(country_meta) = country_meta$standardName
country_meta = country_meta[, c("officialName", "region", "subregion", "subregion2")]
head(country_meta)
```

```
##                               officialName        region          subregion
## India                    Republic of India          Asia      Southern Asia
## China           People's Republic of China          Asia       Eastern Asia
## United States     United States of America North America   Northern America
## Indonesia            Republic of Indonesia          Asia South-Eastern Asia
## Pakistan      Islamic Republic of Pakistan          Asia      Southern Asia
## Nigeria        Federal Republic of Nigeria        Africa     Western Africa
##                       subregion2
## India         South Central Asia
## China                       <NA>
## United States               <NA>
## Indonesia                   <NA>
## Pakistan      South Central Asia
## Nigeria       Sub-Saharan Africa
```

```
df = read.table("num_cite_country_country.tab", quote = "", sep = "\t", header = TRUE)
n_cited = tapply(df[, 3], df[, 1], sum)
n_citing = tapply(df[, 3], df[, 2], sum)


df = read.table("num_cite_country_country.tab", quote = "", sep = "\t", header = TRUE)
df1 = calc_stat(df, all = FALSE); rownames(df1) = paste(df1[, 1], df1[, 2], sep = "-")
df2 = df[df[, 1] != df[, 2], ]
df2 = calc_stat(df2, all = FALSE); rownames(df2) = paste(df2[, 1], df2[, 2], sep = "-")

cn = intersect(rownames(df1), rownames(df2))
df1 = df1[cn, ]
df2 = df2[cn, ]

l = df1$log2_fc- df2$log2_fc < -0.4
plot(df1$log2_fc, df2$log2_fc, cex = ifelse(l, 0.8, 0.2), 
    col = circlize::add_transparency(ifelse(l&grepl("United States", rownames(df1)), "red", 
              ifelse(l&grepl("China", rownames(df1)), "blue", 
                ifelse(l&grepl("Ethiopia", rownames(df1)), "orange", "black"))), 0.25),
    pch = ifelse(df1$country_cited == "United States", 4, 
              ifelse(df1$country_cited == "China", 4,
                 ifelse(df1$country_cited == "Ethiopia", 4, 16))
        ),
    xlab = "log2 fold enrichment / all citations", ylab = "log2 fold enrichment / international citations", main = "compare universe"
)
abline(v = 0, lty = 2, col = "grey")
abline(h = 0, lty = 2, col = "grey")
legend("topleft", cex = 0.7,
    legend = c("from United States", "on United States", "on China", "on Ethiopia", "others"), 
    col = c("red", "red", "blue", "orange", "black"), pch = c(16, 4, 4, 4, 16))
```

```
df = read.table("num_cite_country_country.tab", quote = "", sep = "\t", header = TRUE)
df = df[df[, 1] != df[, 2], ]
df = calc_stat(df, cutoff_total = 10000, cutoff_single = 100, min_country = 24, all = FALSE)
m = xtabs(log2_fc ~ country_cited + country_citing, data = df)
class(m) = "matrix"


all_countries = rownames(m)

m2 = m
m2[m2 < 0] = 0
library(igraph)

set.seed(123)
g = graph_from_adjacency_matrix(m2, mode = "plus", weighted = TRUE)
cm = cluster_louvain(g, weight = E(g)$weight, resolution = 1.2)
communities(cm)
```

```
## $`1`
## [1] "Argentina" "Brazil"    "Chile"     "Colombia"  "Mexico"    "Peru"     
## [7] "Uruguay"  
## 
## $`2`
##  [1] "Australia"      "Austria"        "Belgium"        "Canada"        
##  [5] "Denmark"        "Estonia"        "Finland"        "France"        
##  [9] "Germany"        "Ireland"        "Israel"         "Japan"         
## [13] "Luxembourg"     "Netherlands"    "New Zealand"    "Norway"        
## [17] "Sweden"         "Switzerland"    "United Kingdom" "United States" 
## 
## $`3`
## [1] "Bangladesh"   "Ethiopia"     "Ghana"        "Kenya"        "Nigeria"     
## [6] "South Africa" "Tanzania"     "Uganda"      
## 
## $`4`
##  [1] "Bulgaria"       "Croatia"        "Czech Republic" "Greece"        
##  [5] "Hungary"        "Italy"          "Lithuania"      "Poland"        
##  [9] "Portugal"       "Romania"        "Russia"         "Serbia"        
## [13] "Slovakia"       "Slovenia"       "Spain"         
## 
## $`5`
## [1] "China"       "Hong Kong"   "Indonesia"   "Malaysia"    "Singapore"  
## [6] "South Korea" "Taiwan"      "Thailand"    "Vietnam"    
## 
## $`6`
##  [1] "Egypt"                "India"                "Iran"                
##  [4] "Jordan"               "Kuwait"               "Lebanon"             
##  [7] "Morocco"              "Pakistan"             "Qatar"               
## [10] "Saudi Arabia"         "Tunisia"              "Turkey"              
## [13] "United Arab Emirates"
```

```
mem_by_log2_fc = readRDS("country_groups.rds")
mem_by_log2_fc = cola:::relabel_class(mem_by_log2_fc, mem, return_map = FALSE)
mm = rbind(as.character(mem), as.character(mem_by_log2_fc))
colnames(mm) = names(mem)
Heatmap(mm, name = "classification",
    top_annotation = HeatmapAnnotation(Region = country_meta[rownames(m), "region"], 
                                     col = list(Region = region_color)),
    column_order = order(mem, mem_by_log2_fc, country_meta[rownames(m), "region"]),
    col = structure(names = 1:7, 2:8),
    row_labels = c("universe_international", "universe_all"))
```
